# Safeguarding Genome Integrity: Polo-like kinase Cdc5 and phosphatase Cdc14 Orchestrate Topoisomerase II-Mediated Catenane Resolution in Mitosis

**DOI:** 10.1101/2025.11.06.686928

**Authors:** Lucia F. Massari, Alice Finardi, Clara Visintin, Erika Calabrese, Ambra Dondi, Rosella Visintin

**Author notes:** These authors contributed equally to the work (Lucia F. Massari, Alice Finardi).

## Abstract

Resolution of sister chromatid intertwines in mitosis is vital for safeguarding genome integrity. While cohesin, condensin, the polo-like kinase Cdc5, and the phosphatase Cdc14 have all been implicated in this process, the underlying molecular mechanisms remain elusive. Our study unveils a coordination between spindle elongation and the timely resolution of DNA linkages, orchestrated by Cdc14 and Cdc5. We show that Cdc14 and Cdc5 collaborate to facilitate the resolution of DNA catenanes, the predominant species of DNA intertwines in mitosis, by regulating Topoisomerase II (Top2) function, both through SUMOylation and phosphorylation. Our findings contribute to unraveling a mechanism wherein Cdc5 and Cdc14 work together to ensure the timely removal of all sources of linkages between sister chromatids, thereby preserving the integrity of chromosome segregation and genome stability.

## Introduction

During mitosis, the accurate distribution of the newly replicated genetic material relies on a tightly-regulated process called chromosome segregation. To ensure proper partitioning, the duplicated chromosomes, referred to as sister chromatids, must be paired and recognized as sisters. Cohesin, a ring-shaped protein complex belonging to the structural maintenance of chromosomes (SMC) family, plays a pivotal role in holding sister chromatids together from their synthesis until their separation at anaphase (reviewed in (1)), with cohesin cleavage marking anaphase entry (2, 3).

Besides cohesin, sister chromatids are also connected by DNA linkages called sister chromatid intertwines (SCIs), which arise as byproducts of DNA replication and can be classified as non-replicated DNA segments, recombination intermediates or double-stranded catenanes (reviewed in (4)). To ensure accurate chromosome segregation, all sources of cohesion between sister chromatids must be removed. While most SCIs are resolved in S phase, some persist in mitosis and, in anaphase, appear as threads of DNA spanning between the two segregating DNA masses. Such threads are referred to as anaphase bridges (4). Failure to adequately address these residual linkages can lead to double-stranded DNA breaks during cytokinesis. Even a single unresolved anaphase bridge can cause gross chromosomal rearrangements and genome instability, a hallmark of cancer development (5).

The complete resolution of SCIs is linked to cell cycle events and requires the organization of chromatin into chromosomes, the bipolar attachment of sister chromatids onto the mitotic spindle, and the activity of specialized enzymes. Chromosome formation consists in the non-overlapping compaction of each DNA molecule, which results in the individualization of sister chromatids. Individualization supports the resolution of SCIs by confining their presence to a small region at the interface between the chromatids, thus enhancing the accessibility of the enzymes deputed to their resolution. This process is achieved through the organization of chromatin into loops that are formed by the extruding activity of SMC complexes. In vertebrates the SMC complex carrying out chromosome formation is condensin (6–8), while cohesin, by holding sister chromatids in close proximity, counteracts catenane resolution (9). Indeed, in vertebrates catenanes persist up to metaphase only at the centromere, the only region that still retains cohesin in this stage (10). In the budding yeast *Saccharomyces cerevisiae*, instead, chromosomes are folded mainly by cohesin (11), that has the ability to introduce links both between sister DNA molecules, to achieve cohesion, and within the same molecule, to form loops. Nevertheless, also in budding yeast cohesin counteracts SCI resolution. It has been shown that SCIs can be fully resolved only after cohesin has been cleaved at anaphase onset and sister chromatids are allowed to separate (12, 13).

SCI resolution is facilitated also by the bipolar attachment of sister chromatids onto the spindle, the microtubule structure responsible for pulling them apart. Biorientation and the tension that results from the pulling force exerted by the spindle on centromeres result in the delocalization of condesin from the centromere to chromosome arms, where it can promote SCI resolution (14, 15). It is not known whether anaphase spindle elongation contributes to SCI resolution as well.

Finally, a set of enzymes is dedicated to SCI resolution. Non-replicated DNA patches and recombination intermediates can be cleaved by the mitotic nucleases Yen1 and Mus81-Mms4 (16, 17). Double-stranded catenanes disentanglement relies instead on the action of Type II Topoisomerases (Top2 in budding yeast). Top2 activity is essential for maintaining genome integrity and indeed *top2* mutants are not viable (18–20). Top2 facilitates replication fork progression at termination, facilitates transcription, allows chromatid individualization and compaction and is necessary for catenane resolution (21, 22).

From coordinating the spatial and temporal aspects of key mitotic events to regulating individual enzymatic activities, post-translational modification (PTMs) exert critical roles (23). These include phosphorylation, ubiquitination, and SUMOylation (conjugation with small ubiquitin-like modifier). In budding yeast, two major regulators of mitotic events are the polo-like kinase Cdc5 and the Cdk-counteracting phosphatase Cdc14. Both proteins are integral components of two interconnected signaling networks governing late mitotic events, the FEAR (Cdc14 Early Anaphase Release) and the MEN (Mitotic Exit Network) pathways (reviewed in (24, 25)). The FEAR network facilitates the timely release of Cdc14 from its nucleolar inhibitor Cfi1/Net1 at anaphase onset (26–28). It orchestrates events including sister chromatid separation, nuclear movement, nucleolar segregation (24) and anaphase spindle elongation (29). The MEN pathway instead activates Cdc14 during late anaphase and is crucial for Cdk inactivation, leading to mitotic exit and entry into a new G1 phase (25).

Cdc5 and Cdc14 have also been implicated in SCI resolution. Cdc5 phosphorylates and activates the endonuclease Mus81-Mms4 in G2/M; while in anaphase, Cdc14 activates the endonuclease Yen1 through dephosphorylation (reviewed in (30)). Cdc5 indirectly influences the SUMOylation of Top2 in metaphase by inactivating the SUMO protease Ulp2 (31). It has been suggested that SUMOylation of Top2 directs its recruitment to chromatin (32, 33). Moreover, Cdc5 also promotes condensin activity, the establishment of a bipolar spindle and timely cohesin cleavage.

Although several actors implicated in SCI resolution are known, the intricate molecular mechanisms that govern the disentanglement of DNA during mitosis and their interaction with other mitotic processes remain largely uncharacterized. In this study, we investigate the roles of Cdc5 and Cdc14 in this process. We provide evidence that Cdc5 and Cdc14 orchestrate the timely resolution of the three classes of DNA linkages. We show that during mitosis, Cdc5 and Cdc14 facilitate the resolution of DNA catenanes by influencing Top2 localization, likely by impacting its SUMOylation and phosphorylation status. This function of Cdc5 and Cdc14 operates in parallel with their role in modulating spindle elongation. By coupling sister chromatid intertwine resolution with spindle elongation, these two proteins ensure that sister chromatid segregation occurs together with the removal of all sources of linkages, thereby preserving genomic integrity.

## Materials and methods

### Yeast strains and growth conditions

All yeast strains were W303 (K699) derivatives (Supplementary Table 1). Cell cycle arrest and synchronization followed standard protocols (34). Unless specified, cells were grown at 23 °C in Yeast Extract Peptone (YEP) medium with 2% glucose, raffinose, or galactose. G1, S-phase and metaphase arrests were induced with 5 µg/ml α-factor (Genscript), 10 mg/ml hydroxyurea (Sigma) and 15 µg/ml nocodazole (Sigma, dissolved in DMSO), respectively. To release from the arrests, cells were washed with 10 volumes of fresh medium lacking the pheromone/drug and released into appropriate conditions of medium and temperature (see Fig. legends for details relative to specific experiments). When pertinent, drug(s) were added and restrictive conditions applied. Temperature-sensitive alleles were inactivated at 37 °C. Analog-sensitive *cdc5-as1* and *cdc15-as1* strains were inhibited with 5 µM CMK (custom-made, Accendatech; (35)) or 1NM-PP1 analogue 9 (A603003; Toronto Research Chemicals). The *CDC20-AID* allele was inactivated with 500 µM auxin (I5148;Sigma), and *MET-CDC20* was repressed with 8 mM methionine.

### Indirect in situ immunofluorescence

Immunofluorescence (IF) was performed as described in (36). Briefly, 1 ml of culture at OD₆₀₀ = 0.4–0.6 was fixed overnight in 3.7% formaldehyde/0.1 M potassium phosphate buffer (pH 6.4) at 4 °C. Cells were washed three times in 0.1 M potassium phosphate buffer and once in sorbitol buffer (1.2 M sorbitol, 0.1M K_2_HPO_4,_ 0.33 M citric acid, pH 5.9), then digested with 0.1 mg/ml Zymolyase 100T for ∼30 min at 30 °C. Spheroplasts were washed, adhered to poly-L-lysine–coated slides, and sequentially treated with cold methanol (−20 °C, 3 min) and acetone (−20 °C, 10 s). Speroplasts were incubated 90 min with primary antibodies diluted in PBS/BSA (1% BSA, 0.04 M K₂HPO₄, 0.01 M KH₂PO₄, 0.15 M NaCl, 0.1% NaN₃); washed; incubated 90 min with fluorophore-conjugated secondary antibodies; washed and covered with DAPI mounting solution (90% glycerol, 10% KPBS, 100 mg p-phenylenediamine, 5 μg DAPI).

Primary antibodies were rat anti–α-tubulin YOL34 (1:100, AbD Serotec), mouse anti-Nop1 (1:2000, EnCore), and mouse anti-PK SV5-Pk1 (1:500, AbD Serotec). Secondary antibodies were donkey anti-rat (FITC 1:100, Jackson; Cy3 1:100, Jackson) and donkey anti-mouse (FITC 1:100, Jackson; Cy3 1:500, Jackson). Slides were sealed with nail polish and analyzed by fluorescence microscopy.

### Microscopy and image analysis

Cell cycle progression was monitored by categorizing ≥100 cells *per* sample as interphase, metaphase, or anaphase based on spindle morphology. For spindle length and nuclear morphology, images were acquired on an upright Olympus AX70 Provis microscope with a 100×/1.40 oil UPlanSApo∞/0.17/FN 26.5 objective and a Photometrics CoolSnap 12-bit B&W camera using MetaMorph 7.5.6.0 software (MDS Analytical Technologies), and analyzed with Fiji. For anaphase bridge and spindle measurements, images were collected on a DeltaVision Elite deconvolution system (Applied Precision) equipped with an Olympus IX71 inverted microscope, UPlanSApo 100× oil objective (NA 1.4), and a CoolSnap HQ2 CCD camera. Z-stacks (12 × 0.5 µm) were acquired with FITC, TRITC, and DAPI filters, deconvoluted with SoftWoRx, and analyzed with Fiji to assess nuclear and nucleolar morphology and measure spindle length. Anaphase cells were scored for DNA and/or nucleolar bridges using DAPI and Nop1 staining. Intensity thresholds were defined in wild-type controls lacking bridges (4000 a.u. for DAPI, 5200 a.u. for Nop1). Cells exceeding the DAPI threshold were classified as DAPI-bridge positive, those exceeding the Nop1 threshold as nucleolar-bridge positive, and cells above both thresholds as double-bridge positive. Cells below both thresholds were scored negative.

For Top2 localization, cells stained for tubulin, Top2-PK, and DNA (DAPI) were imaged on a Nikon Eclipse Ti2 microscope with a spinning-disk X-Light V3 module (CrestOptics), solid-state lasers (Lumencor Celesta), a Kinetix sCMOS camera, and a 100×/1.49 NA oil objective. Z-stacks (25 planes, 0.2 µm spacing) were acquired and deconvoluted using the Blind algorithm in NIS. Briefly, images were resampled to obtain isotropic voxels, ensuring geometric accuracy for 3-D morphometry. The DAPI channel was then denoised (Gaussian blur with σ = 2) and segmented by four-level Multi-Otsu thresholding. The resulting segmented nuclei were filtered to based on their volume to remove debris and aggregates. The nuclear masks were then expanded to enclose the Top2 signal (5 px corresponding to 325 nm). From each expanded mask, the mean and standard deviation of the Top2 signal intensity were calculated. The coefficient of variation for each mask was calculated by dividing the standard deviation by the mean. At least 250 cells *per* strain were analyzed. Statistical significance was assessed using the Kruskal-Wallis test followed by Mann-Whitney U test for post-hoc pairwise comparisons.

### Live-cell imaging

Cells were grown in SCD medium supplemented with 2% adenine and released from G1 arrest into Y04C microfluidic plates (CellASIC). Imaging was performed on a DeltaVision microscope as previously described. To avoid UV-induced DNA damage checkpoint activation, live imaging began once cells reached metaphase, identified by spindle morphology and nuclear positioning. For spindle elongation (Fig. 1; Supplementary Figs. 2– 3) and nuclear/nucleolar segregation (Fig. 1; Supplementary Figs. 1–2), 8 z-stacks (0.6 µm step) were acquired every 3 min using FITC and Cherry filters with a 100× oil objective. For GFP-dot separation (Fig. 2; Supplementary Figs. 4–5), 15 z-stacks (0.4 µm step) were acquired every 3 min under the same conditions. Reference images were collected in DIC. Stacks were deconvoluted with SoftWoRx software, and spindle length and chromosome locus distances were measured in x, y, z using Fiji with a custom plug-in (Spindle Z manual tracker). Representative images are shown as maximum-intensity projections.

**Fig. 1:**
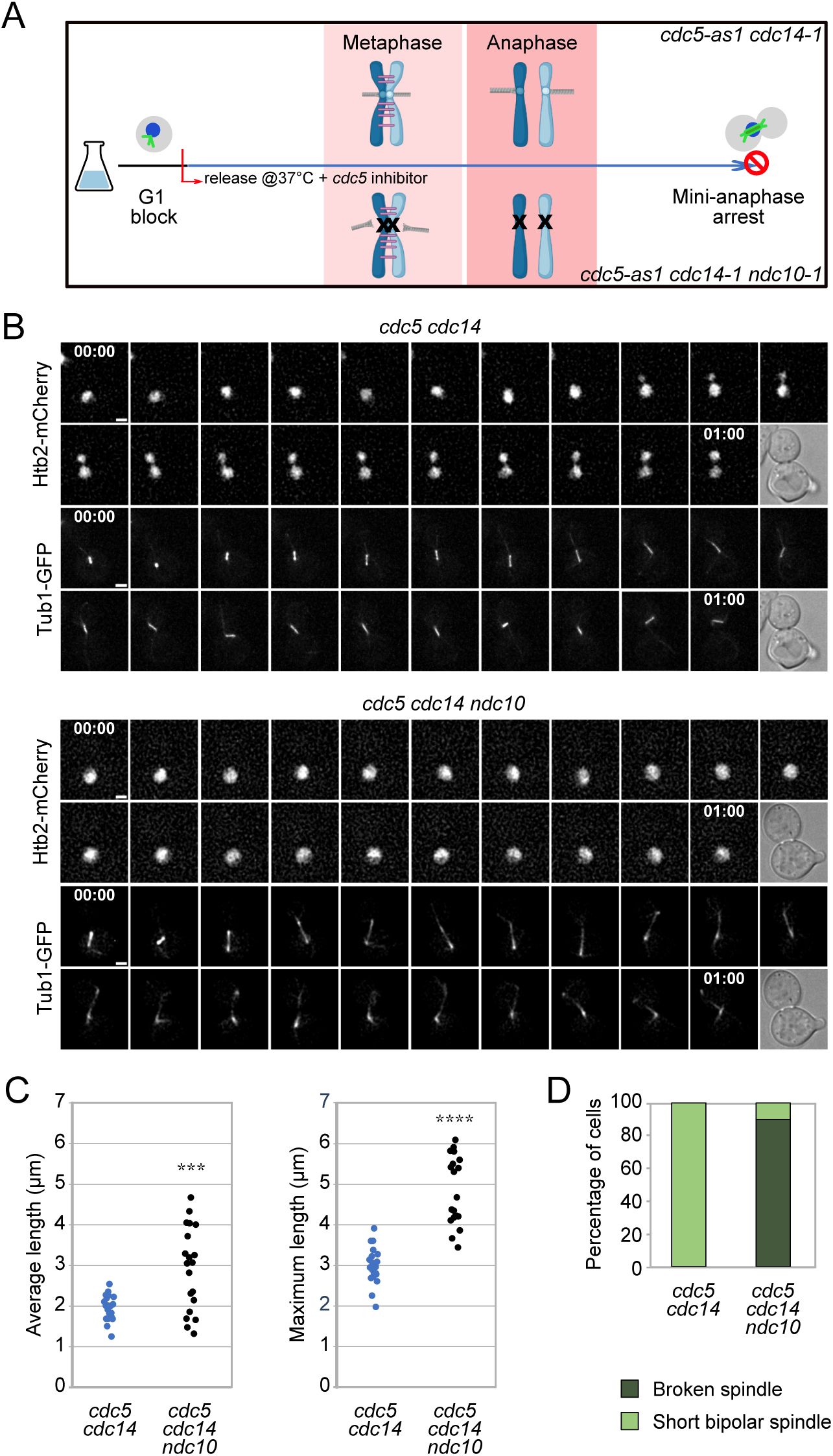
*cdc5 cdc14* cells are defective in sister chromatid separation. *cdc5-as1 cdc14-1* (Ry6591) and *cdc5-as1 cdc14-1 ndc10-1* (Ry6589) cells carrying a *HTB2-Cherry* and a *GFP-TUB1* fusion were synchronously released from a G1 block in conditions restrictive for *cdc5*, *cdc14*, and *ndc10*. A schematic of the rationale and experimental setup is shown (**A**). Cells were imaged starting from metaphase every 3 min for 3 hours. n=20 cells for each strain. Representative images of the first hour of imaging are shown (**B**). Scale bars = 2 μm. Average and maximum spindle length for each cell analyzed are shown (**C)**. NOTE: each dot represents one cell. The average spindle length in *cdc5 cdc14 ndc10* cells (mean = 3.0 μm, S.D. = 1) was higher than in *cdc5 cdc14* cells (mean = 2.0 μm, S.D. =0.3). *** = P<0.001. The maximum spindle length in *cdc5 cdc14 ndc10* (mean = 4.8 μm, S.D. = 0.8) was higher than in *cdc5 cdc14* cells (mean = 3.0 μm, S.D. = 0.5). **** = P<0.0001. Percentage of cells that either maintained a short bipolar spindle or broke the spindle during the time-lapse (**D**).

**Fig. 2:**
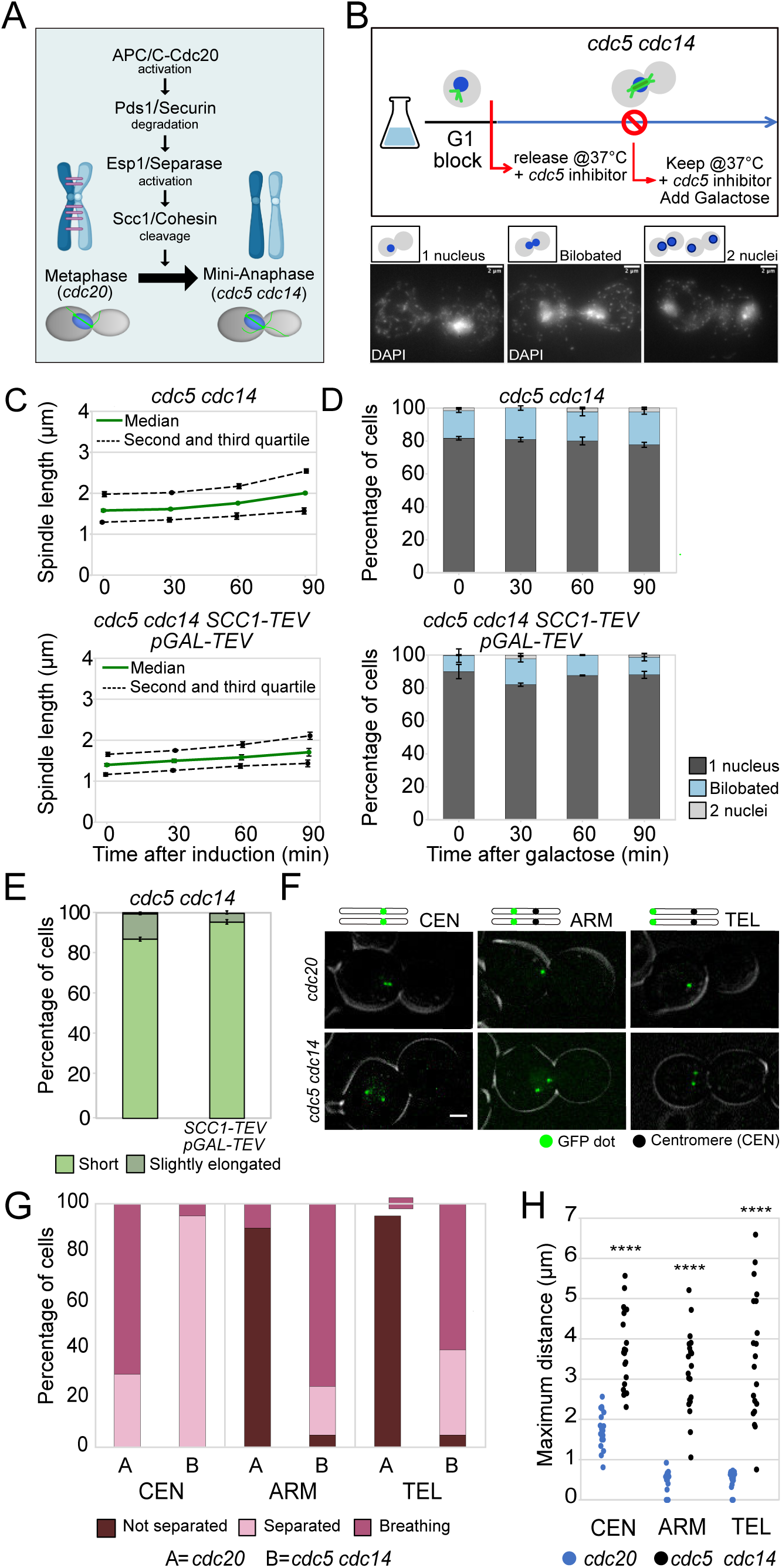
Cohesin does not contribute to the sister chromatid separation defect of *cdc5 cdc14* mutants. Schematic representation of the signaling cascade leading to cohesin cleavage, the critical step for anaphase entry (**A**). *cdc5-as1 cdc14-1* (Ry1602) and *cdc5-as1 cdc14-1 SCC1-TEV GAL-TEV* (Ry2795) cells were synchronously released from a G1 block in YPR in conditions restrictive for *cdc5* and *cdc14*. At the *cdc5 cdc14* arrest (3h30 after release), galactose was added to induce TEV protease expression. Samples were collected at the indicated time points after induction and analyzed through indirect IF (anti-Tub1, DAPI) to monitor spindle length and nuclear morphology. The schematic diagram of the experimental setup and images representative of nuclear morphology categories are shown. Scale bars = 2 μm (**B**). Spindle length: Median (solid green line), second and third quartile (dotted lines) and s.e.m. (error bars) are shown (**C**). Nuclear and spindle morphology: Mean and s.e.m. (error bars) are shown (**D-E**). Three independent experiments are shown, n=100 cells counted for each time point in each experiment. *CDC20-AID* and *cdc5-as1 cdc14-1* cells carrying a *CENXV-GFP (*Ry7519 and Ry5815), a *HIS3-GFP* (Ry7411 and Ry5824) or a *TELXV-GFP* (Ry7481 and Ry7098) fusion were synchronously released from a G1 block into conditions restrictive for *cdc20*, *cdc5* and *cdc14*. Cells were imaged starting from metaphase every 3 minutes for 3 hours. n=20 cells were analyzed in each condition. Representative images are shown. Scale bars = 2 μm (**F**). Distance between GFP dots. According to the distance between GFP dots throughout the time-lapse, cells were assigned to the following categories: Cells in which the distance between dots consistently remained below 2 μm = Not separated; Cells in which the distance between dots exceeded 2 μm, but subsequently returned to zero = Breathing; Cells in which the distance between dots exceeded 2 μm and never went back to zero = Separated (**G**). Maximum distance between GFP dots for each cell analyzed. The distance was higher in *cdc5 cdc14* cells compared to *cdc20* cells at every locus analyzed (**H**). **** = P<0.0001.

### Immunoblot analysis

Cells were treated with cold 5% TCA for 10 min, pelleted, and washed with acetone. Pellets were resuspended in 50 mM Tris-HCl (pH 7.5), 1 mM EDTA, 1 mM p-nitrophenyl phosphate, 50 mM DTT, 1 mM PMSF, and 2 μg/ml pepstatin, lysed with glass beads, and boiled in 1× SDS sample buffer.

Proteins were detected with the following primary antibodies: mouse anti-PK (SV5-Pk1, AbD Serotec; 1:5000) for Top2-9PK, rabbit anti-Smt3 (gift from Dr. Branzei; 1:2000) for SUMOylated species, and mouse anti-Pgk1 (A-6457, Molecular Probes; 1:5000) as a loading control. Secondary antibodies were HRP-conjugated goat anti-rabbit IgG (Bio-Rad, 170-6515; 1:5000) and HRP-conjugated goat anti-mouse IgG (Bio-Rad, 170-6516; 1:10000). Signals were visualized by chemiluminescence (ECL; GE Healthcare).

Phos-tag gels were prepared with 40 μM Phos-tag AAL-107 (Fujifilm Wako) and 10 mM MnCl₂ in 6% acrylamide gels.

### Immunoprecipitation of Top2

Cells were lysed as in (31). After resuspending the pellet in the SDS-containing solution, the lysate was boiled and clarified by centrifugation. The supernatant was diluted 1:10 with NP40 buffer and incubated with SV5-Pk1 monoclonal mouse anti-PK antibody, used at 1:200 dilution, and Pierce protein-G agarose beads (Thermo Scientific). The beads were washed with NP40. The beads were resuspended in 3x SDS sample buffer to elute the proteins.

### Co-immunoprecipitation

A 50 ml culture was grown to OD₆₀₀ ≈ 0.5, harvested at 3, 000 rpm for 5 min at 4 °C, and resuspended in 150 µl NP-40 buffer with protease and phosphatase inhibitors. Cells were lysed using FastPrep (4 × 45 s, maximum speed, 4 °C), and lysates clarified at 13, 000 rpm for 10 min. Protein concentration was determined by Bradford, and 500 µg extract (50 µl) was used per IP, adjusted to equal volumes with NP-40 buffer. For immunoprecipitation, 10 µl anti-V5 magnetic beads (Top2-PK, SAE0203, Sigma) or 15 µl Pierce™ anti-HA magnetic beads (Cdc14-HA or Cdc5-HA, 88837, ThermoFisher) were added and rotated 2 h at 4 °C. Beads were washed five times with NP-40 buffer and eluted in 30 µl 3× Laemmli buffer, boiled 5 min, centrifuged, and supernatants analyzed by SDS–PAGE.

### Calibrated Chromatin ImmunoPrecipitation and sequencing (ChIP-Seq)

ChIP-seq was performed as described in of (37) with modifications from (38) (39). For each condition, 200 ml of *S. cerevisiae* cultures (*TOP2-PK9*) arrested in metaphase were fixed in 20 ml fixing solution (6 ml 37% formaldehyde, 143 mM NaCl, 1.43 mM EDTA, 71.4 mM HEPES-KOH pH 7.5) for 80 min at room temperature, quenched with 0.125 M glycine for 5 min, pelleted, and washed twice in 10 ml of ice-cold TBS (20 mM Tris-HCl pH 7.5, 150 mM NaCl) and once in 10 ml ice-cold 1xFA lysis buffer (50 mM HEPES-KOH pH 7.5, 140 mM NaCl, 1 mM EDTA, 1% Triton X-100, 0.1% Na deoxycholate) added with 0.1% SDS. Pellets were snap-frozen in liquid nitrogen and stored at –80 °C.

For calibration, *S. cerevisiae* pellets were mixed 1:1 with *S. pombe* (*rad21-V5:KANMX6*) pellets, prepared as follow. *S. pombe* cells were grown in YES media (0.5% yeast extract, 3% glucose, 2% agar 225 mg/l each of adenine, histidine, leucin, uracil, and lysine hydrochloride) at 30°C until reaching OD_600_ = 0.4-0.6. The culture was fixed, quenched, harvested and washed like the *S.cerevisiae* one. *S. cerevisiae* and *S. pombe* cell pellets were thawed on ice and each *S. cerevisiae* pellet was combined with one *S. pombe* pellet in 400 μl 1xFA* lysis buffer (FA buffer added with 1× Roche EDTA-free protease inhibitors and 1 mM PMSF) with 0.5% SDS. Cells were lysed with silica beads (FastPrep) and chromatin was sheared by sonication (Bioruptor Plus, 2 × 20 cycles, 30 s ON/OFF, high power). Soluble, sheared, chromatin was collected after centrifugation, pooled, and split into input and IP samples. Inputs were diluted 1:50 in TE and stored overnight at 4 °C. For IPs, 1 ml of extract was incubated either with 15 µl protein A Dynabeads (Invitrogen) or 10 µl V5 antibody (Bio-Rad) overnight at 4 °C. Beads were washed as follow: twice in cold 1ml 1xFA lysis buffer/0.1%SDS; twice in cold 1ml 1xFA lysis buffer/0.1%SDS+360mM NaCl (total: 500mM NaCl); twice in 1ml cold 10mMTris–HCl, pH 8.0 250mMLiCl 0.5% NP-40 0.5% Na-deoxycholate 1mM EDTA; once in 1ml cold TE pH 8. Inputs and IPs were decrosslinked overnight with proteinase K at 65 °C, purified with the Wizard kit (Promega), and stored in 35 µl water at –20 °C.

DNA libraries were prepared from 2 ng DNA as in from (38) DNA ends were blunted and phosphorylated (NEB Quick Blunting kit), <100 bp fragments removed (AMPure XP beads), and dA tails added (NEB Klenow exo-). Each sample was then ligated with a different NEXTflex-6 DNA Barcode (PerkinElmer) using the Quick Ligation kit (NEB), unligated adapters removed and libraries PCR-amplified (Phusion polymerase, NEB) using primers specific for NextFlex adapters (5’-AATGATACGGCGACCACCGAGATCTACAC; 5’-CAAGCAGAAGACGGCATACGAGAT). Fragments (150–300 bp) were size-selected with AMPure XP, quantified by Qubit, and quality-checked by Bioanalyzer (Agilent) using the 2100 Bioanalyzer High Sensitivity DNA kit (Agilent, Santa Clara, CA).

INPUT and IP libraries for each sample were pooled at a 15:85 ratio and sequenced on an Illumina MiniSeq with a 150-cycle high-output kit.

Data analysis was performed as in (37). Reads were mapped to both the *S. pombe* calibration genome and the *S. cerevisiae* sacCer3 genome. Occupancy ratios (OR = (INPUT_cal_ × IP_exp_) / (INPUT_exp_ × IP_cal_)) were calculated and used to normalize read counts.

Region-specific plots were generated with the Spark package (https://github.com/harbourlab/SparK). Mean calibrated ChIP-seq plots (all chromosome pileups) were generated and visualised with computeMatrix and plotProfile in deepTools (ref: https://academic.oup.com/nar/article/44/W1/W160/2499308). In plots showing 3Kb flanking the centromere, reads were binned at 50bp, in plots showing 80Kb flanking the centromere, at 500bp. In plots showing the chromosome arm from cen to tel, each arm was averaged to 10Kb and binned to 50bp. Centromere coordinates used are listed in Supplementary Table 2.

### Statistical analysis

Depending on the experiment and as indicated in the corresponding Fig. legends, P values were determined by unpaired or paired Student’s t-test after arcsine transformation as appropriate for the data obtained from a count and expressed as percentages. A P value of less than 0.05 was considered statistically significant (∗ = P < 0.05; ∗∗ = P < 0.01, ∗∗∗ = P < 0.001 and **** = P < 0.0001). Averages ± s.e.m. (standard error of the mean) of three independent experiments are reported.

### Repeatability of the experiments

Each experiment in the manuscript has been repeated at least three independent times. The results were highly reproducible.

## Results

### Mini-anaphase arrested *cdc5 cdc14* cells are defective in sister chromatid separation

To investigate how DNA linkages are removed during mitosis, we analyzed budding yeast cells lacking both the Polo-like kinase Cdc5 (*cdc5-as1,* henceforth *cdc5*) (35) and the Cdk-counteracting phosphatase Cdc14 (*cdc14-1,* henceforth *cdc14*) (40). The *cdc5-as1 cdc14-1* double mutant strain (henceforth *cdc5 cdc14*) arrests in “mini-anaphase”, a state characterized by a short, stable bipolar mitotic spindle and unseparated nuclei despite cohesin cleavage, due to defective spindle elongation (29), and, potentially, to linkages between sister chromatids. To further characterize the mini-anaphase arrest of *cdc5 cdc14* cells, we performed live-cell imaging with Htb2-mCherry and GFP-Tub1 to capture chromosome segregation dynamics under restrictive conditions (Fig. 1A). In stark contrast to the results obtained through fixed-sample indirect immunofluorescence (IF), which showed a single undivided nuclear mass (29), live imaging revealed highly dynamic nuclei that often appeared bilobated (Fig. 1B). Such transient morphology, overlooked in population-based assays, likely reflects nuclei initiating separation and reuniting due to spindle defects, or being compressed while passing through the bud neck. The bilobated phenotype can also originate from the nucleus and nucleolus residing in two different cellular compartments, as already documented for mutants associated with the G2/M DNA damage checkpoint and with the FEAR network (41, 42). Indeed, the nucleolus, which contains ribosomal DNA (rDNA) repeats, is inadequately stained by DAPI, but it can be effectively labeled with Htb2-Cherry. The latter enabling clear visualization of both the nucleus and the nucleolus. To discriminate among these possibilities, we examined *cdc5 cdc14* cells carrying Htb2-Cherry and Cfi1-GFP fusions, the latter marking the nucleolus. In all cells examined, the nucleus migrated into the daughter cell, while the nucleolus remained in the mother (Supplementary Fig. 1A and B). During its passage through the bud-neck, the nucleus appeared bilobated due to compression (Supplementary Fig. 1A). Therefore, such morphology reflects both nuclear compression at the bud neck and the spatial separation of nucleus and nucleolus, consistent with known roles of Cdc14 and Cdc5 in nucleolar segregation (43–45). Importantly, this phenotype does not represent chromosome segregation, confirming that *cdc5 cdc14* cells are defective in this process. Previous data suggested that the chromosome segregation defect of *cdc5 cdc14* cells is independent of spindle elongation (29). Disrupting chromosome attachment to the spindle by inactivating the kinetochore protein Ndc10, *ndc10-1* (Fig. 1A) (46), altered spindle morphology in double mutant cells. Unlike *cdc5 cdc14* cells, which formed short and stable spindles, *ndc10 cdc5 cdc14* mutants developed fragile spindles that often broke during attempted elongation (29). To further characterize this phenotype, we analyzed nuclear morphology and spindle dynamics in *ndc10*, *cdc5 cdc14*, and *cdc5 cdc14 ndc10* cells. In line with the absence of microtubule attachment to chromosomes, both *ndc10* and *cdc5 cdc14 ndc10* cells displayed a single nucleus that remained in the mother cell throughout the experiment (Supplementary Fig. 2A and Fig. 1B). Spindle measurements supported the persistence of residual cohesion in *cdc5 cdc14* cells. In this strain, the spindles remained stable and short, with an average length ranging from 1.2 to 2.8 μm (mean of 2 μm) and a maximum length ranging from 2 to 4 μm (mean of 3.0 μm) (Fig. 1B, C and Supplementary Fig. 3A). Instead, in *cdc5 cdc14 ndc10* cells the spindles were longer, reaching a maximum length of 6 μm (mean of 4.8 μm) and an average length ranging from 1.2 to 4.8 μm (mean of 3 μm) (Fig. 1B, C and Supplementary Fig. 3B). Furthermore, spindles in this mutant displayed increased fragility, collapsing in 90% of the analyzed cells during the experiment (Fig. 1D, Supplementary Fig. 3B). Consistent with the spindle elongation defect observed in *cdc5 cdc14* mutants, *cdc5 cdc14 ndc10* cells failed to achieve full spindle elongation, unlike *ndc10* cells, which successfully elongated their spindles up to 14 μm (mean of 9 μm), and completed the cell cycle (Supplementary Fig. 2B). Together, these results indicate that unresolved cohesion between sister chromatids persists in *cdc5 cdc14* cells. This residual linkage counteracts the pulling forces of a compromised spindle, preventing full chromatid separation.

### Linkages other than cohesin contribute to the sister chromatid separation defect of *cdc5 cdc14* cells

The residual cohesion observed in *cdc5 cdc14* cells could arise either from uncleaved cohesin or from sister chromatid intertwines. Although separase Esp1 is active and cleaves bulk cohesin in these cells (Fig. 2A) (29), it remained possible that a fraction of cohesive cohesin escaped cleavage. To address this, we applied two complementary strategies: ectopic cohesin cleavage and direct analysis of cohesion at specific chromosomal loci.

For ectopic cleavage, we overexpressed the Tobacco Etch Virus (TEV) protease (47) under the galactose-inducible *GAL1-10* promoter (*GAL-TEV)* in strains carrying a TEV-cleavable allele of the *SCC1* subunit of cohesin (*scc1Δscc1-TEV268,* henceforth *SCC1-TEV*) (3). If residual cohesin were responsible for cohesion, TEV induction in *cdc5 cdc14* cells should mimic the phenotype of *cdc5 cdc14 ndc10* cells, where chromatids no longer restrain spindle elongation (Fig. 1). After synchronization and TEV induction at the *cdc5 cdc14* block (Fig. 2B and Supplementary Fig. 4A ), we measured spindle length and categorized cells into three groups based on nuclear morphology - with a single nucleus, a separating nuclear mass (bilobated), or with two divided nuclei (Fig. 2B) - in *cdc5 cdc14* and *cdc5 cdc14 SCC1-TEV GAL-TEV* cells. Consistent with our previous report (29), *cdc5 cdc14* cells maintained short and stable spindles (1-2.5 μm) (Fig. 2C) and largely undivided nuclei (Fig. 2D). The shorter spindle length in this experiment, as compared to the one in Fig. 1, is attributed to the use of a less rich media, containing raffinose instead of glucose. Remarkably, ectopic cohesin cleavage (Supplementary Fig. 4A) did not elicit spindle elongation nor induce nuclear division in *cdc5 cdc14 SCC1-TEV GAL-TEV* cells (Fig. 2C and D) with spindles remaining short and stable (Fig. 2E). Thus, anaphase cohesion in *cdc5 cdc14* cells does not originate from residual cohesin.

We next monitored cohesion at individual chromosomal loci using the TetR-GFP/*tetO* system (48). Distinct chromosomal regions exhibit different behaviors depending if they are bound by cohesin or not. When bound by cohesin (*e.g.* in metaphase), sister telomeres and chromosome arms remain tightly associated, and, when labelled, appear as one fluorescent dot. Conversely, at the centromeres the tension exerted by the mitotic spindle counteracts the cohesive function of cohesin, leading to a phenomenon known as "centromere breathing”, where sister centromeres continuously separate (two dots) and rejoin (one dot) over distances shorter than 2 μm (49–51). We examined a centromere (*CENXV*), a chromosome arm (*HIS3*), and a telomere (*TELXVr*), all on the long arm of chromosome XV ((52) and Fig. 2F), comparing *cdc20* metaphase-arrested cells (with intact cohesin) to *cdc5 cdc14* cells (Fig. 2A). Chromosome XV was chosen due to its lack of specialized sequences and its length (it is one of the longest yeast chromosomes), which could exacerbate a sister chromatid separation defect. The *cdc20* metaphase arrest was obtained by depleting Cdc20, the activating subunit of the anaphase-promoting complex/cyclosome (APC/C) (Fig. 2A) with the *CDC20-AID* allele, that allows conditional degradation of Cdc20 upon auxin treatment (53). In the absence of Cdc20, the APC/C cannot be activated and cells remain arrested in metaphase with uncleaved cohesin and high Cdk activity (54). In designing the experiment, we planned to visualize spindle pole bodies (SPBs) to monitor mitosis and reference GFP dot positioning. However, integrating a fluorescently tagged SPB component into the *cdc5 cdc14* background either led to lethality or caused synthetic interactions that disrupted normal cell cycle progression and SPB dynamics, preventing our analysis. To distinguish coordinated breathing (elastic motion) from the random movement of two dots, for a preliminary analysis, we used the *SPC42*-mScarlet allele - this allele significantly slows down cell cycle progression, but preserves spindle pole body dynamics (Supplementary Fig. 4B). After confirming that we were not observing random movements, we investigated chromosome behavior in the absence of a SPB reference. As expected (49, 50), *cdc20* cell showed centromeric breathing (two dots oscillating within ∼2 µm) (Fig. 2F, 2G and Supplementary Fig. 5A), while arms and telomeres were largely undivided (Fig. 2F, Fig. 2G and Supplementary Fig. 5B, 5C). In contrast, *cdc5 cdc14* cells displayed extensive separation: centromeres split up to 5–6 µm without rejoining (Fig. 2F, 2G and Supplementary Fig. 5A), and arms and telomeres displayed significant separation (Fig. 2F, 2G and Supplementary Fig. 5B, 5C), reaching distances up to 4-5 μm and 5-6 μm, respectively (Fig. 2H). These patterns are incompatible with stable cohesin-mediated cohesion. Notably, some *cdc5 cdc14* cells showed a breathing-like rejoining of arms and telomeres even after separation >2 µm (Supplementary Fig. 5B, 5C), suggesting either stochastic movement, chromatin reorganization, or persistence of non-cohesin linkages. If the latter is true, then these linkages may be more prominent on chromosome arms than on telomeres.

Together these results indicate that in *cdc5 cdc14* cells: (i) centromeres and pericentromeric regions lack cohesin or other linkages; (ii) most cohesin is cleaved and removed genome-wide; and (iii) chromosome arms and telomeres likely retain SCI-derived linkages.

### Cell lacking Cdc5 and Cdc14 activities accumulate mitotic SCIs

To determine whether cohesin-independent cohesion in *cdc5 cdc14* cells derives from sister chromatid intertwines, we examined the presence of DNA anaphase bridges (4). To visualize anaphase bridges in *cdc5 cdc14* cells we forced spindle elongation, hence chromosome segregation, by overexpressing the Kinesin 5 motor protein Cin8 (55). The phospho-null constitutively active *cin8-3A* allele (56, 57) was previously shown to rescue spindle elongation in these cells (29). Here, we expressed *CIN8* under the galactose-inducible *GAL1-10* promoter, allowing induction after cells had reached the *cdc5 cdc14* arrest (Fig. 3A). As readouts, we analysed spindle length, nuclear morphology, and anaphase bridges by indirect immunofluorescence (IF). Since Cdc5 and Cdc14 are required for the segregation of the nucleolus (43–45), to exclude the possibility that bridges reflected only rDNA segregation defects, we stained the nucleolar marker Nop1 (58). Threads decorated only by Nop1 were classified as nucleolar bridges. Instead, DAPI-stained threads were scored as anaphase/DAPI bridges, regardless of the presence of Nop1 (Supplementary Fig. 6A).

**Fig. 3:**
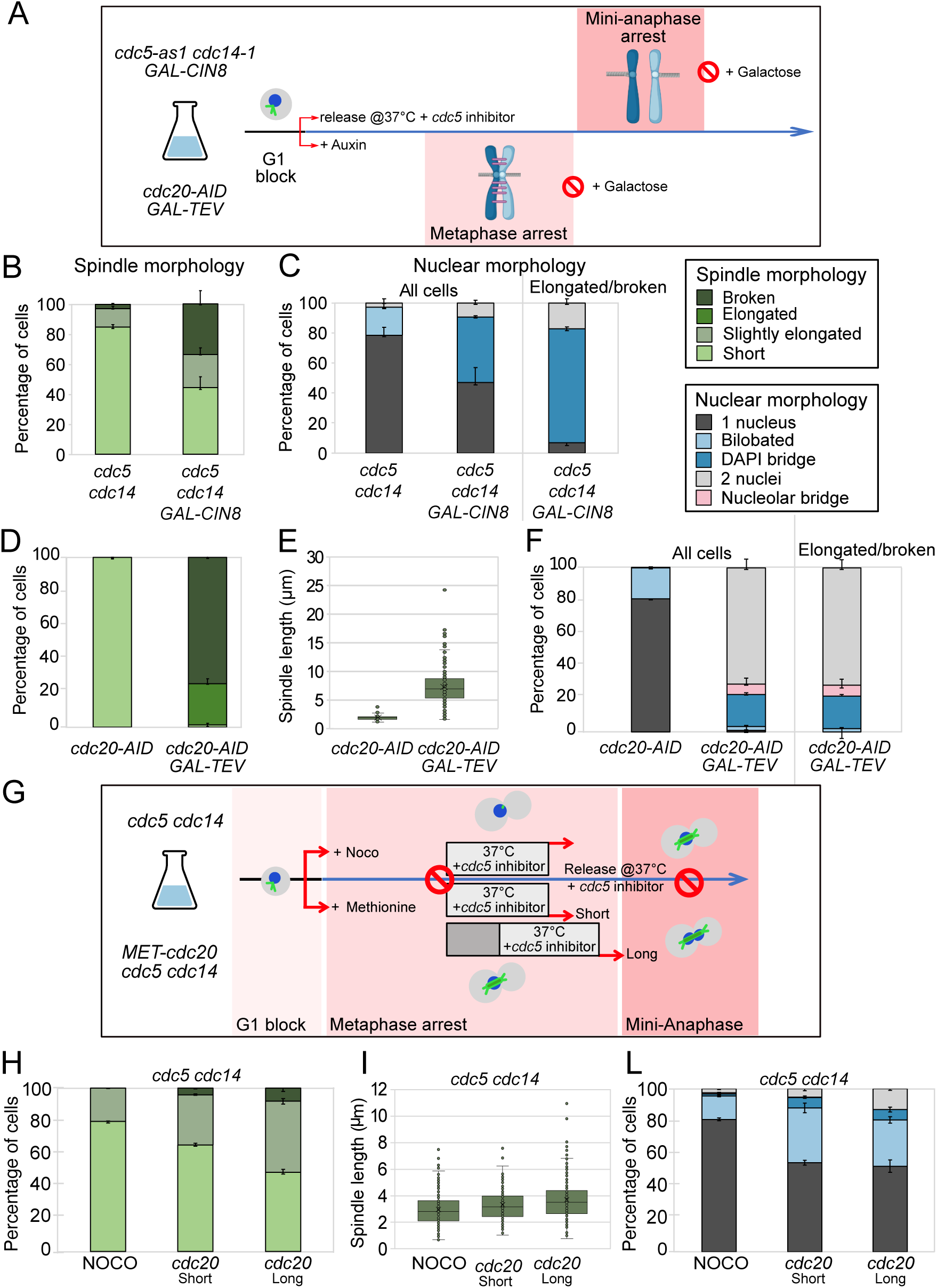
Cdc5 and Cdc14 promote the mitotic resolution of sister chromatid intertwines. Forcing spindle elongation in *cdc5 cdc14* cells originates anaphase bridges. *cdc5-as1 cdc14-*1 (Ry1602) and *cdc5-as1 cdc14-1 GAL-CIN8* (Ry5956) cells were synchronously released from a G1 block in YPR in conditions restrictive for *cdc5* and *cdc14*. At the *cdc5 cdc14* arrest (3h30 after release), galactose was added to induce Cin8 overexpression. A schematic of the rationale and experimental setup is shown (**A**). Samples were collected 120 min after induction and analyzed through indirect IF (anti-Tub1, DAPI, anti-Nop1) to monitor spindle (**B**) and nuclear and nucleolar morphologies (**C**) respectively. Ectopic cohesin cleavage in metaphase-arrested cells leads to spindle elongation without forming anaphase bridges. *CDC20-AID*(Ry4852) and *CDC20-AIDSCC1-TEV GAL-TEV* (Ry4931) cells were synchronously released from a G1 block in YPR in conditions restrictive for *cdc20*. At the metaphase arrest (3h after release), galactose was added to induce TEV protease expression. Samples were collected 120 min after induction and analyzed through indirect IF (anti-Tub1, DAPI) to monitor spindle morphology (**D**), spindle lenght (**E**) and nuclear morphology (**F**). Cdc14 and/or Cdc5 activity requires tension establishment. *cdc5-as1 cdc14-1* (Ry1602) and *cdc5-as1 cdc14-1 MET-CDC20* (Ry3203) cells were synchronously released from a G1 block into a metaphase arrest by adding the depolymerizing drug, nocodazole (NOCO), and methionine (*cdc20*), respectively. When the arrest was complete Cdc5 and Cdc14 were inhibited for 45 minutes before release maintaining restrictive conditions (NOCO and *cdc20* short). Half culture of cells carrying the *MET-CDC20* allele/ were kept arrested for an additional 30 minutes (*cdc20* long) before inactivating Cdc5 and Cdc14 and next treated as above. A schematic of the rationale and experimental setup is shown (**G**). Samples were collected at the indicated time after release and analyzed through indirect IF (anti-Tub1, DAPI) to monitor spindle morphology (**H**), spindle length (**I**) and nuclear morphology (**L**). Mean and s.e.m. (error bars) from three independent experiments are shown. n=200 cells were analyzed for each condition in each experiment.

Following galactose induction, spindles in *cdc5 cdc14* cells remained short and stable, whereas in *cdc5 cdc14 GAL-CIN8* cells they elongated and broke in approximately 60% of cases (Fig. 3B and Supplementary Fig. 6B). Nuclear morphology was unchanged in *cdc5 cdc14* cells but separation occurred concurrently with spindle elongation in *cdc5 cdc14 GAL-CIN8* cells (Fig. 3C). Notably, 73% of cells with elongated or broken spindles displayed DAPI bridges, confirming the persistence of DNA linkages in *cdc5 cdc14* cells at the terminal arrest (Fig. 3C and Supplementary Fig. 6A). Nucleoli remained undivided in both strains and did not contribute to bridging.

### Cdc5 and/or Cdc14 are required for the mitotic resolution of SCIs

The persistence of DNA linkages in *cdc5 cdc14* cells raised the question of whether these are a normal feature of mini-anaphase or a direct consequence of lacking Cdc5 and/or Cdc14 activity. Since mini-anaphase cannot be assayed independently of the *cdc5 cdc14* mutant, at least in the W303 background, we examined wild-type cells arrested in metaphase and forced into segregation by ectopic cohesin cleavage (3). If spindle elongation alone allows segregation without anaphase bridges, SCI resolution must require Cdc5 and/or Cdc14. To test this, we arrested *cdc20* and *cdc20 SCC1-TEV GAL-TEV* cells in metaphase, induced TEV protease, and analyzed samples 120 minutes later (Fig. 3A). Because Cdc14 is activated sequentially by FEAR and MEN, MEN requires spindle pole body entry into the daughter cell (25), and in *cdc20*-arrested cells the nucleus migrates into the daughter compartment (59), we also included *cdc20 cdc15* mutants to exclude MEN contribution (reviewed in (60)). Following cohesin cleavage (Supplementary Fig. 7A) and consistent with prior studies (3), spindle elongated and collapsed in the majority (∼99%) of *cdc20 GAL-TEV* and *cdc20 cdc15 GAL-TEV* cells (Fig. 3D and Supplementary Fig. 7B), with spindles reaching an average length of 6-7 μm (Fig. 3E and Supplementary Fig. 7C). Remarkably, only 27% of the cells with elongated and broken spindles displayed DAPI-positive bridges, compared with 73% in *cdc5 cdc14 GAL-CIN8* cells (Fig. 3F and Supplementary Fig. 7D vs Fig. 3C). Thus, the metaphase biochemical environment can resolve DNA intertwines once cohesin is cleaved and spindles elongate, whereas mini-anaphase without Cdc5 and Cdc14 cannot. To exclude incomplete cohesin cleavage as the cause of the segregation defect in *cdc5 cdc14* cells, we compared Scc1 processing in *cdc20 SCC1TEV GAL-TEV* and *cdc5 cdc14 SCC1TEV GAL-TEV* strains, analyzing samples side by side. In both cases, full-length Scc1 was nearly undetectable 120 minutes after galactose induction (Supplementary Fig. 7E). Yet, only *cdc20* cells completed chromatid separation and bridge resolution, while *cdc5 cdc14* cells remained unsegregated despite similar residual Scc1 levels (Fig. 3D-F vs Fig. 2C-D). These results indicate that the segregation failure in *cdc5 cdc14* cells does not stem from incomplete cohesin cleavage but rather from persistent DNA intertwines. Together, these findings demonstrate that full SCI resolution requires Cdc5 and/or early Cdc14 activity, independently of MEN function.

### Mitotic disentanglement of DNA linkages requires Cdc5, Cdc14 and an active bipolar spindle during both metaphase and anaphase

In previous experiments, Cdc5 and Cdc14 were inactivated from G1 onward. To pinpoint when these proteins promote SCI resolution, we tested their inactivation specifically in metaphase and simultaneously asked whether their role is mediated through the control of spindle dynamics (29), (61). Cells were arrested in metaphase using an allele of *CDC20* under the control of the methionine-repressible promoter (*MET-CDC20*), the sole mutant allele of *CDC20* viable in the *cdc5 cdc14* double mutant background (29), either with or without a bipolar spindle (the latter by nocodazole treatment - NOCO). Once arrests were established, Cdc5 and Cdc14 were inactivated for 45 min before release into anaphase (Fig. 3G) in restrictive conditions. Spindle and nuclear morphologies were evaluated 120 minutes post metaphase release (Fig. 3H-L).

Both conditions produced a mini-anaphase–like phenotype. In nocodazole-treated cells, ∼80% showed a single nucleus and a short/stable spindle, whereas cells released from a *cdc20* arrest (referred to as *cdc20* short arrest) displayed a phenotype resembling *cdc5 cdc14 ndc10* mutants: ∼40% formed slightly elongated spindles (mean 3.8 µm) and ∼50% segregated their DNA (Fig. 3H–L). These different outcomes are most likely explained by differences in the tension exerted on chromatin, rather than by spindle checkpoint activation in nocodazole-treated cells (62), since Securin degradation and Scc1 cleavage confirmed SAC inactivation after nocodazole release (Supplementary Fig. 7F). To assess whether metaphase activity alone is sufficient, we extended the *cdc20* arrest for 30 min before inactivating Cdc5 and Cdc14 (Fig. 3G). This “long arrest” improved outcomes—60% of cells elongated spindles to ∼4 µm and segregated DNA (Fig. 3H–L; Supplementary Fig. 7G)— but still failed to mimic *cdc5 cdc14 ndc10* mutants (∼ 80% of cells with broken spindles), suggesting that SCI resolution requires Cdc5 and Cdc14 activity also from anaphase onward.

We next compared anaphase phenotypes of *cdc5*, *cdc14*, and *cdc15* single mutants. Inactivation of either Cdc5 or Cdc14 increased anaphase bridges relative to *cdc15* cells, but with distinct profiles: *cdc14* mutants exhibited a lower fraction of cells with anaphase bridges but retained nucleolar bridges, while *cdc5* mutants accumulated chromosomal bridges, particularly at early anaphase when spindles were short (Fig. 4A, B and 4E). Quantification of bridges relative to spindle length across mutants and synchronized wild-type cells revealed a correlation between bridge resolution and spindle length (Fig. 4C, D). However, both *cdc5* and *cdc14* mutants retained bridges even when spindles exceeded 8 µm, whereas *cdc15* and wild-type cells did not (Fig. 4C, E and Supplementary Fig. 8A). Thus, spindle elongation facilitates SCI removal but is insufficient without Cdc5 or Cdc14. This aligns with the findings that Cin8 overexpression in *cdc5 cdc14* cells elongates spindles but fails to resolve bridges.

**Fig. 4.**
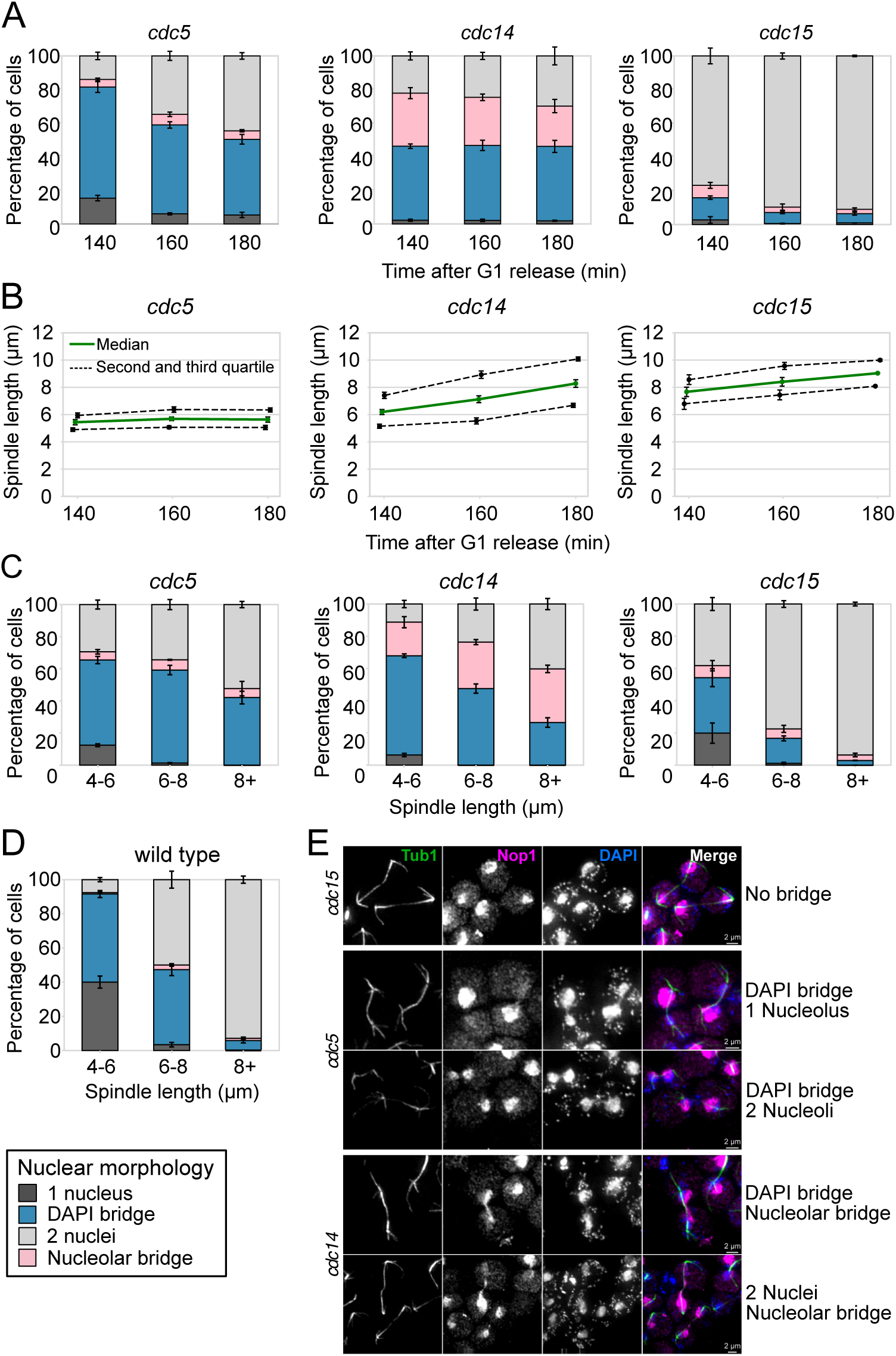
Anaphase bridges resolution requires spindle elongation and Cdc5 and Cdc14 activities. Wild-type (Ry1), *cdc5-as1* (Ry2446), *cdc14-1* (Ry1573) and *cdc15-as1* (Ry1112) cells were synchronously released from a G1 block in conditions restrictive for *cdc5*, *cdc14* and *cdc15*. Samples were collected at the indicated times after release and analyzed through indirect IF (DAPI, anti-Nop1, anti-tubulin) to monitor nuclear and nucleolar morphologies (**A**) and spindle length (**B**). n= 100 cells counted for each time point in each experiment. Spindles were measured and cells were assigned to the indicated categories according to nuclear and nucleolar morphology. n=100 anaphase cells (spindle > 4 μm) were analyzed at each time point. For wild-type cells we chose samples from 80 min to 120 min after release (**D**), while for *cdc5*, *cdc14*, and *cdc15* cells we analyzed samples 140 min, 160 min, and 180 min after release (**C**). Representative images for each strain are shown. Scale bars = 2 μm (**E**). Mean and s.e.m. (error bars) from three independent experiments are shown.

Together, these results demonstrate that SCI resolution requires Cdc5 and Cdc14 both in metaphase and anaphase. In anaphase, they act in parallel with regulating spindle elongation, with Cdc5 primarily promoting chromosomal bridge resolution and Cdc14 nucleolar segregation.

### DNA catenanes hinder sister chromatid separation in *cdc5 cdc14* mutants

To understand how Cdc5 and Cdc14 promote the resolution of DNA linkages, we sought to identify which SCI species persist in their absence. Sister chromatid can be linked by non-replicated DNA, recombination intermediates, or double-stranded catenanes (4). Because replication of difficult regions can extend into anaphase (63), and Cdc5 and Cdc14 regulate nucleases acting on replication and recombination intermediates, we first tested whether these structures underlie the segregation defect. Both replication and recombination intermediates can be resolved by Yen1. We therefore introduced a constitutively active allele, *YEN1^ON^* (64), into *cdc5 cdc14* cells. *YEN1^ON^* modestly improved spindle elongation and nuclear division (Fig. 5A-C), similar to the effect of overexpressing wild-type Yen1 (Supplementary Fig. 8B and C). In *cdc5 cdc14 GAL-CIN8* cells, *YEN1^ON^*allowed further spindle elongation and nuclear separation, but reduced anaphase bridges only slightly (Fig. 5D-F). As ∼65% of cells still retained bridges, recombination and replication intermediates likely contribute to - but do not primarily account for - the persistence of DNA linkages in this mutant.

**Fig. 5:**
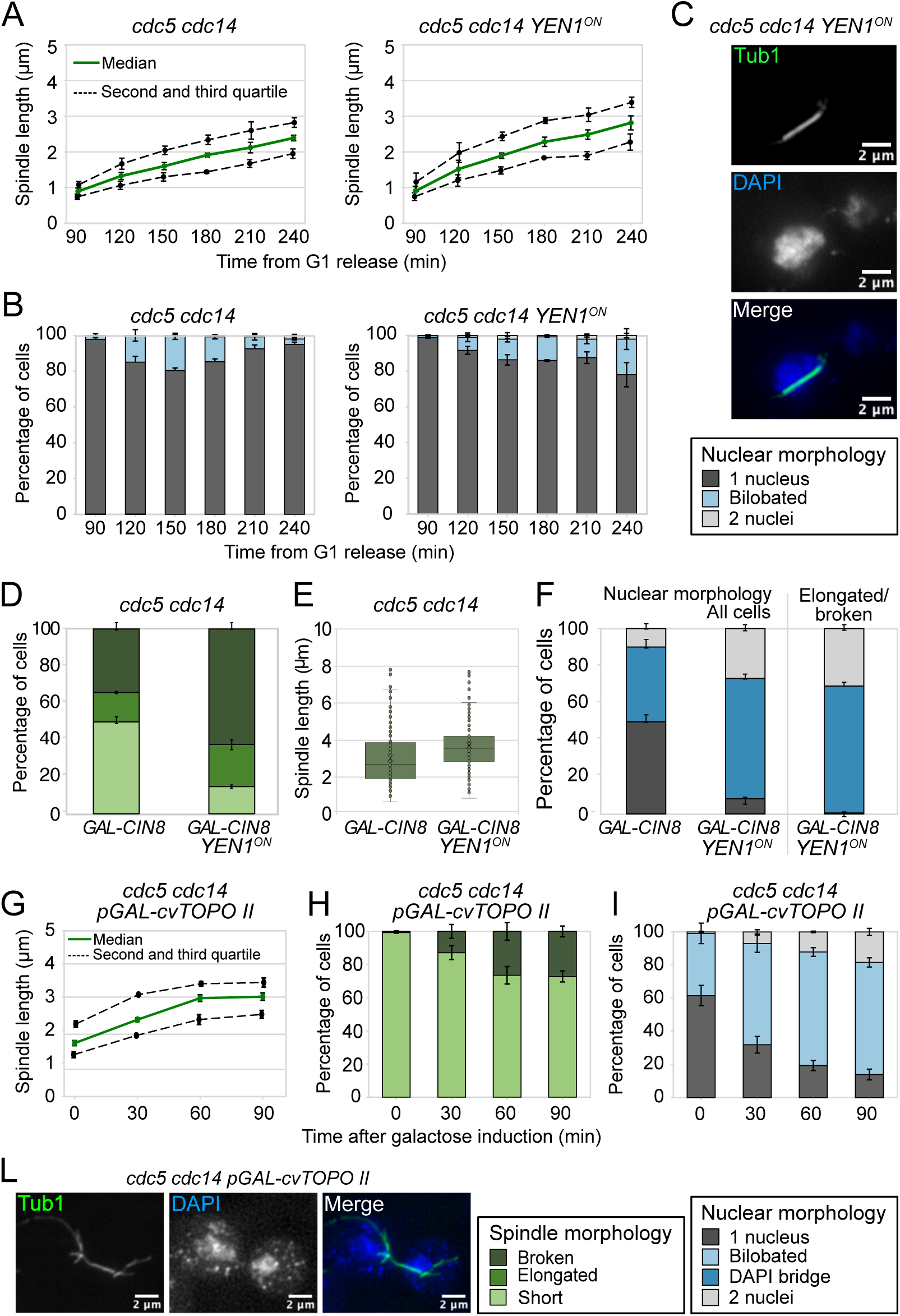
Cdc5 and Cdc14 promote catenane resolution in mitosis. *cdc5-as1 cdc14-1* (Ry1602) cells and *cdc5-as1 cdc14-1 YEN1^ON^-MYC* (Ry5272) cells were synchronously released from a G1 block in conditions restrictive for *cdc5* and *cdc14*. Samples were collected at the indicated time points after release and analyzed through indirect IF (anti-Tub1, DAPI) to monitor spindle length (**A**) and nuclear morphology (**B**). Representative images for the *cdc5-as1 cdc14-1 YEN1^ON^-MYC* are shown. Scale bars = 2 μm (**C**). *cdc5-as1 cdc14-1 GAL-CIN8* (Ry5956) and *cdc5-as1 cdc14-1 GAL-CIN8 YEN1^ON^-MYC* (Ry7220) cells were synchronously released from a G1 block in YPR in conditions restrictive for *cdc5* and *cdc14*. At the *cdc5 cdc14* arrest (3h30 after release), galactose was added to induce Cin8 overexpression (schematic diagram in Fig. 3A) Samples were collected at the indicated time to monitor spindle morphology (**D**), spindle lenght (**E**) and nuclear morphology (**F**). *cdc5-as1 cdc14-1* (Ry1602) and *cdc5-as1 cdc14-1 GAL-cvTOPOII* (Ry5156) cells were synchronously released from a G1 block in YPR in conditions restrictive for *cdc5* and *cdc14*. At the terminal arrest (3h30 after release), galactose was added to induce cv-TopoII overexpression. At the indicated time points spindle length (**G**) and morphology (**H**) and nuclear morphology (**I**) were assessed. Representative images for the *cdc5-as1 cdc14-1 GAL-cvTOPOII* are shown. Scale bars = 2 μm (**L**). Three independent experiments are shown. n=100 cells counted at each time point in each experiment. Median (solid line), second and third quartile (dotted lines) and s.e.m. (error bars) are shown (**A**, **G**). Mean and s.e.m. (error bars) are shown for the remaining graphs.

We next examined the contribution of DNA catenanes, whose resolution requires type II topoisomerases (Top2 in *S. cerevisiae*). Overexpression of yeast Top2 only marginally improved spindle and nuclear phenotypes of *cdc5 cdc14* cells (Supplementary Fig. 9A-C). By contrast, overexpression of the viral type II topoisomerase from *Paramecium bursaria Chlorella* virus (cv-Topo II), which lacks the C-terminal regulatory domain (65) nearly fully rescued nuclear division and produced elongated but fragile spindles (Fig. 5G–L), similar to those in the *cdc5 cdc14 ndc10* mutant. Notably, cv-Topo II had no effect on *cdc20* metaphase-arrested cells, where cohesin remains uncleaved (Supplementary Fig. 9D, E), indicating that its rescue activity depends on the absence of cohesin.

These findings strongly suggest that unresolved DNA catenanes are the major form of SCI in the *cdc5 cdc14* mutant and that they are sufficient to prevent chromatid separation and spindle elongation in these cells.

### Cdc5 and Cdc14 regulate Top2 recruitment to chromatin and nuclear localization

The complete resolution of DNA catenanes during mitosis requires the coordinated action of a bipolar spindle, condensin, and Topoisomerase II. Given that *cdc5 cdc14* mutants are defective in both spindle elongation (29), and topoisomerase function, we asked whether impaired condensin activity also contributes to their terminal phenotype (15, 43, 44). Because condensin cannot easily be reconstituted *in vivo* and overexpressed, we tested its contribution indirectly. If condensin were inactive in *cdc5 cdc14* cells, the presence of the *smc2-8* loss-of-function allele of condensin should not alter the ability of cv-Topo II to rescue the segregation defect in these cells. Instead, cv-Topo II failed to rescue the SCI resolution defect of *cdc5 cdc14 smc2* cells (Supplementary Fig. 9F, G), indicating that condensin retains sufficient function in the double mutant.

We therefore next investigated how Cdc5 and Cdc14 influence Top2 function. Because Top2 activity relies on its localization within the nucleus and recruitment to chromatin (15, 32, 66), we tested whether these processes were disrupted in *cdc5 cdc14* cells by analyzing Top2 distribution through calibrated ChIP-seq and fluorescence imaging.

We examined Top2 localization with ChIP-seq in *cdc5 cdc14* cells at their terminal arrest. As controls we used a condition in which Top2 localization is known (wild-type cells arrested in metaphase without a bipolar spindle - NOCO) and an intermediate condition in which the other player in catenane resolution, condensin, already displays significant changes (wild-type cells with a bipolar spindle – *cdc20* arrest). Previous studies showed that the tension exerted by the bipolar attachment of sister centromeres onto the metaphase spindle is sufficient to delocalize condensin from the centromere to chromosome arms and that Top2 levels increase on chromosome arms upon release from metaphase (15, 67). As before, we arrested cells in metaphase by inactivation of Cdc20 (*CDC20-AID*) and disrupted the spindle by treatment with nocodazole. Our ChIP-seq results indicate that Top2 is enriched at the centromere in the absence of a bipolar spindle, as shown both on an individual chromosome (full chromsomome III in Fig. 6A, zoom-in on the pericentromere in Fig. 6B) and in a pile-up plot averaging all 16 yeast chromosomes, centered on the centromere (Fig. 6C). This enrichment diminishes with the establishment of tension (Fig. 6A-C), similarly to what happens to condensin (15). However, we did not see a corresponding increase of Top2 levels along chromosome arms, neither looking at individual chromosomes (Fig. 6A), nor by averaging all chromosomes in the 160Kb flanking the centromere (Fig. 6D), nor by averaging all chromosome arms from centromere to telomere (Fig. 6E). Strikingly, *cdc5 cdc14* cells displayed markedly reduced Top2 occupancy at both centromeres and chromosome arms (Fig. 6), suggesting a defect in Top2 recruitment to DNA.

**Fig. 6:**
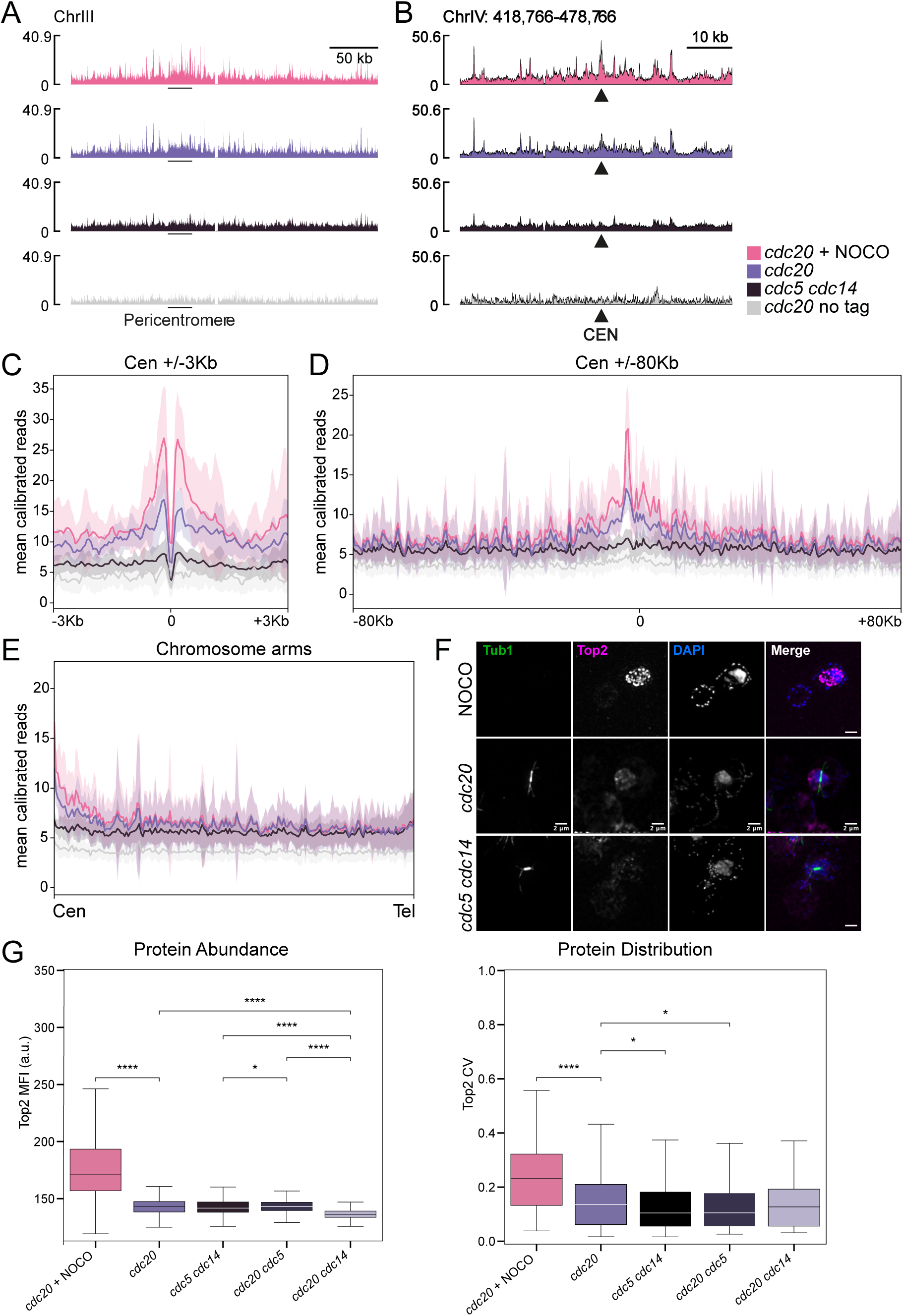
Cdc5 and Cdc14 recruit Top2 to chromatin. (**A-D**) Calibrated Top2 ChIP-seq in *CDC20-AID osTIR* (Ry4852) and *CDC20-AID osTIR TOP2-PK9* (Ry8315) cells arrested in metaphase with and without a spindle and in arrested *cdc5-as1 cdc14-1 TOP2-PK9* cells (Ry7998). Cultures were arrested in G1 and released in presence of auxin (*cdc20, cdc20* NOCO and *cdc20* no tag), nocodazole (*cdc20* NOCO) or DMSO (*cdc20*, *cdc20* no tag and *cdc5 cdc14*) or in restrictive conditions for *cdc5-as1* (*cdc5 cdc14*) or for *cdc14* (37°, all cultures). Cultures were harvested when they reached the metaphase (2.30h after G1 release, *cdc20*, *cdc20* NOCO and *cdc20* no tag) or mini-anaphase arrest (3h after release, *cdc5 cdc14*). Top2 ChIP track along chromosome III. The position of the pericentromere is shown (**A**). Top2 ChIP-seq track in the 60Kb surrounding Centromere IV. The position of centromere IV is shown (CEN) (**B**). Pileup of all 16 centromeres and flanking 3 Kb regions (**C**). Pileup of all 16 centromeres and flanking 80Kb regions. 80kb is the length of the shortest chromosome arm (**D**). Pile-up of all chromosome arms, scaled and oriented from centromere to telomere (**E**). Line: mean calibrated reads, shade: standard deviation (**CDE**). (**F-G**) *CDC20-AID* (Ry8315), *CDC20-AID cdc5-as1* (Ry8785), *CDC20-AID cdc14-1* (Ry8782) and *cdc5-as1 cdc14-1* (Ry7998) cells carrying a *TOP2-9PK* fusion cells treated as above were collected at the terminal arrest for IF against Top2. Representative images of *cdc20* arrest +/- NOCO and *cdc5 cdc14* cells obtained by the path stacks, Z-project, STD projection in Fiji are shown in (**F**) (**G**) Quantification of Top2 abundance and distribution in expanded nuclear regions: left panel shows the mean nuclear Top2 fluorescence intensity (MFI, a.u., proxy for Top2 abundance) and the right panel the coefficient of variation (CV, proxy for Top2 distribution) of voxel-level Top2 signal within each nucleus. Data are presented as box plots with median, quartiles, and whiskers indicating the range. Statistical significance was determined by Kruskal-Wallis test followed by Mann-Whitney U test for post-hoc pairwise comparisons. *P < 0.05, ****P < 0.0001. For each strain, at least 250 cells were analyzed.

Imaging of Top2-PK confirmed these findings. In a metaphase arrest in the absence of bipolar spindle (nocodazole), Top2 formed bright nuclear foci, while in the *cdc20* arrest the overall signal intensity and localization pattern diminished (Fig. 6F, G and Supplementary Tables 3 and 4 for statistics). In *cdc5 cdc14* cells, overall Top2 levels were comparable to *cdc20* cells, but its localization became diffuse, consistent with ChIP-seq. Analysis of *cdc20 cdc5* and *cdc20 cdc14* mutants revealed distinct contributions: loss of Cdc5 largely recapitulated the double mutant phenotype, whereas loss of Cdc14 caused a milder defect in chromatin association but markedly reduced overall Top2 levels, leading to diminished nuclear localization (Fig. 6G and Supplementary Tables 3 and 4 for statistics). Together, these findings indicate that Cdc5 and Cdc14 cooperate to ensure proper Top2 loading onto chromatin and its nuclear distribution during metaphase, providing a mechanistic basis for the defective catenane resolution observed in *cdc5 cdc14* cells.

### Cdc5-dependent mitotic SUMOylation has minimal impact on catenane resolution

To explore how this regulation occurs, we next asked whether Cdc5 and Cdc14 control Top2 localization through post-translational modifications, focusing on SUMOylation—a process that may depend on Cdc5’s established role in inhibiting the SUMO protease Ulp2 (31). Top2 was immunoprecipitated from metaphase-arrested *cdc20*, *cdc20 cdc5*, and *cdc20 cdc14* cells, as well as from *cdc5 cdc14* cells arrested in mini-anaphase. SUMOylation was detected with an anti-Smt3 (yeast SUMO) antibody. Top2–Smt3 bands were observed in *cdc20* and *cdc20 cdc14* cells but not in *cdc20 cdc5* and *cdc5 cdc14* cells, indicating that Cdc5, but not Cdc14, is required for Top2 SUMOylation (Fig. 7A-B and Supplementary Fig. 10A). Consistently, Cdc5 overexpression during an S-phase arrest induced with hydroxyurea treatment—when endogenous Cdc5 is low phase (68, 69) and its activity further inhibited by the HU treatment triggering the DNA damage checkpoint (70–72)— ectopically enhanced Top2 SUMOylation (Fig. 7C), confirming Cdc5’s role in this modification (31–33). To test whether defective SUMOylation accounts for the *cdc5 cdc14* segregation defect, we analyzed *cdc14*, *cdc15*, and *cdc5* mutant cells expressing a non-SUMOylatable Top2 allele (*top2-SNM*) (73). In all cases, *top2-SNM* did not significantly increase anaphase bridges (Supplementary Fig. 10B, C), suggesting that loss of Top2 SUMOylation alone is not sufficient to explain the phenotype.

**Fig. 7:**
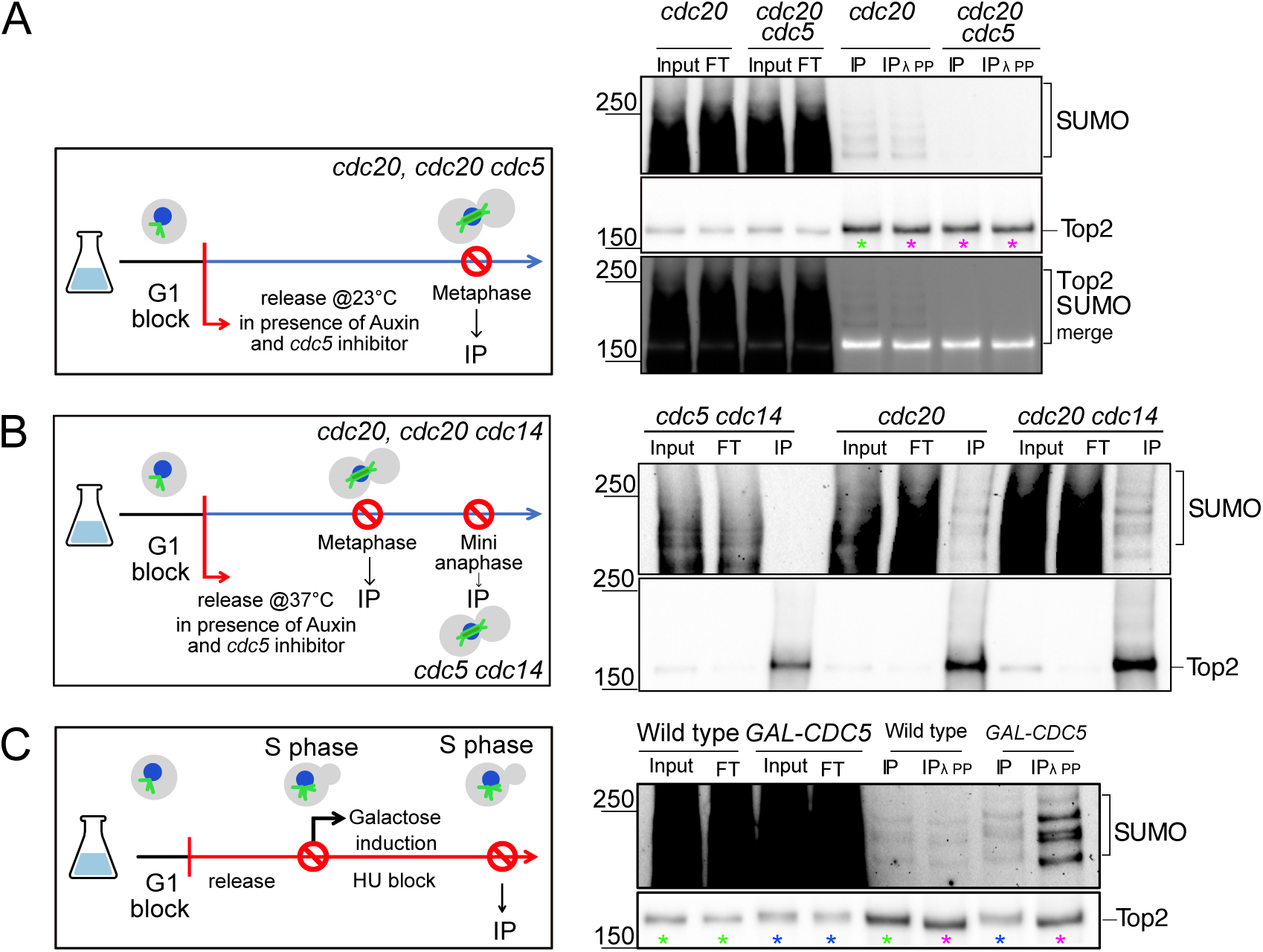
Cdc5 and Cdc14 regulate Top2 function via mechanisms beyond sumoylation. (**A**-**B**) *CDC20-AID* (Ry8315), *CDC20-AID cdc5-as1* (Ry8785), *CDC20-AID cdc14-1* (Ry8782) and *cdc5-as1 cdc14-1* (Ry7998) cells carrying a *TOP2-9PK* fusion were synchronously released from a G1 block in conditions restrictive for the proteins of interest. Samples for Top2 immunoprecipitation were collected at the terminal arrests. Schematic representation of the experimental design and immunostaining against Top2 and Smt3 (SUMO) are shown. (**C**) Wild-type (Ry7921) and *GAL-CDC5-3MYC* (Ry9325) cells carrying a *TOP2-9PK* fusion were arrested in S phase with hydroxyurea in YPR for 3h. At the arrest, galactose was added to induce Cdc5 overexpression. 2h after the induction a sample was taken for Top2 immunoprecipitation. Schematic representation of the experimental design and immunostaining against Top2 and Smt3 (SUMO) are shown. (**A**-**C**) Input, FT, flowthrough; IP, immunoprecipitate; IP+λPP, immunoprecipitate treated with λ phosphatase (PP).

We next asked whether Cdc5’s regulation of sister chromatid intertwines (SCIs) could involve multiple Ulp2 substrates (74). If so, Ulp2 overexpression would be expected to phenocopy *cdc5* defects. After confirming that Ulp2 overexpression reduced global SUMOylation (Supplementary Fig. 10D), we synchronized *cdc5, cdc15*, and *cdc15 GAL-ULP2* cells in nocodazole to accumulate unresolved DNA catenanes (14, 67, 75–77). Upon arrest, Ulp2 expression was induced for 1 hour before nocodazole removal (Supplementary Fig. 10E). In *cdc15* mutants, Ulp2 overexpression delayed both spindle elongation and bridge resolution after release (Supplementary Fig. 10E; F).

Nonetheless, segregation eventually proceeded, and the defects were less severe than those observed in *cdc5* mutants (Supplementary Fig. 10F). These findings indicate that while Cdc5 regulation of Ulp2 and Top2 SUMOylation contributes to SCI resolution, SUMOylation alone does not explain the severe defects of *cdc5 cdc14* cells, implying that additional Cdc5-dependent mechanisms are involved.

### Cdc5 and Cdc14 regulate Top2 phosphorylation

In addition to SUMOylation, Top2 is also regulated by phosphorylation, although its role during mitosis remains less well defined. Given the kinase and phosphatase activities of Cdc5 and Cdc14, we investigated whether they modulate Top2 phosphorylation. Treating immunoprecipitated Top2 from *cdc20*-arrested metaphase cells with λ-phosphatase did not alter its SUMOylation pattern but generated faster-migrating bands on SDS–PAGE—visible for both Top2 and its SUMOylated form (Fig. 7A, magenta vs green asterisks)—mirroring the mobility observed in *cdc5* mutants, indicating phosphorylation control (Fig. 7A, magenta asterisks). Consistently, Cdc5 overexpression in hydroxyurea-treated cells induced a pronounced mobility upshift of bulk Top2 (Fig. 7C, blue *vs.* green asterisk) which was abolished by λ-phosphatase despite SUMOylation remaining intact (Fig. 7C, magenta asterisk), and the appearance of slow mobility forms (Supplementary Fig. 11A),, confirming that Cdc5 promotes Top2 phosphorylation.

We next examined whether Cdc14 contributes to regulating Top2 phosphorylation. In metaphase-arrested *cdc14* cells, Top2 migrated as a slower, higher band compared to G1 (Fig. 8A, blue *vs*. magenta asterisk), which shifted downward upon Cdc14 overexpression, indicating that Cdc14 can dephosphorylate Top2. Whether Cdc14 overexpression shifts Top2 mobility completely to the G1 pattern—mirrored by the *cdc20 cdc5* cells—or to an intermediate state is difficult to determine with certainty (Fig. 8A, green *vs.* blue and magenta asterisks). Notably, the G1-like pattern observed in *cdc20 cdc5* cells remained unchanged upon Cdc14 overexpression, suggesting that Cdc14 might already possess some activity during metaphase.

**Fig. 8:**
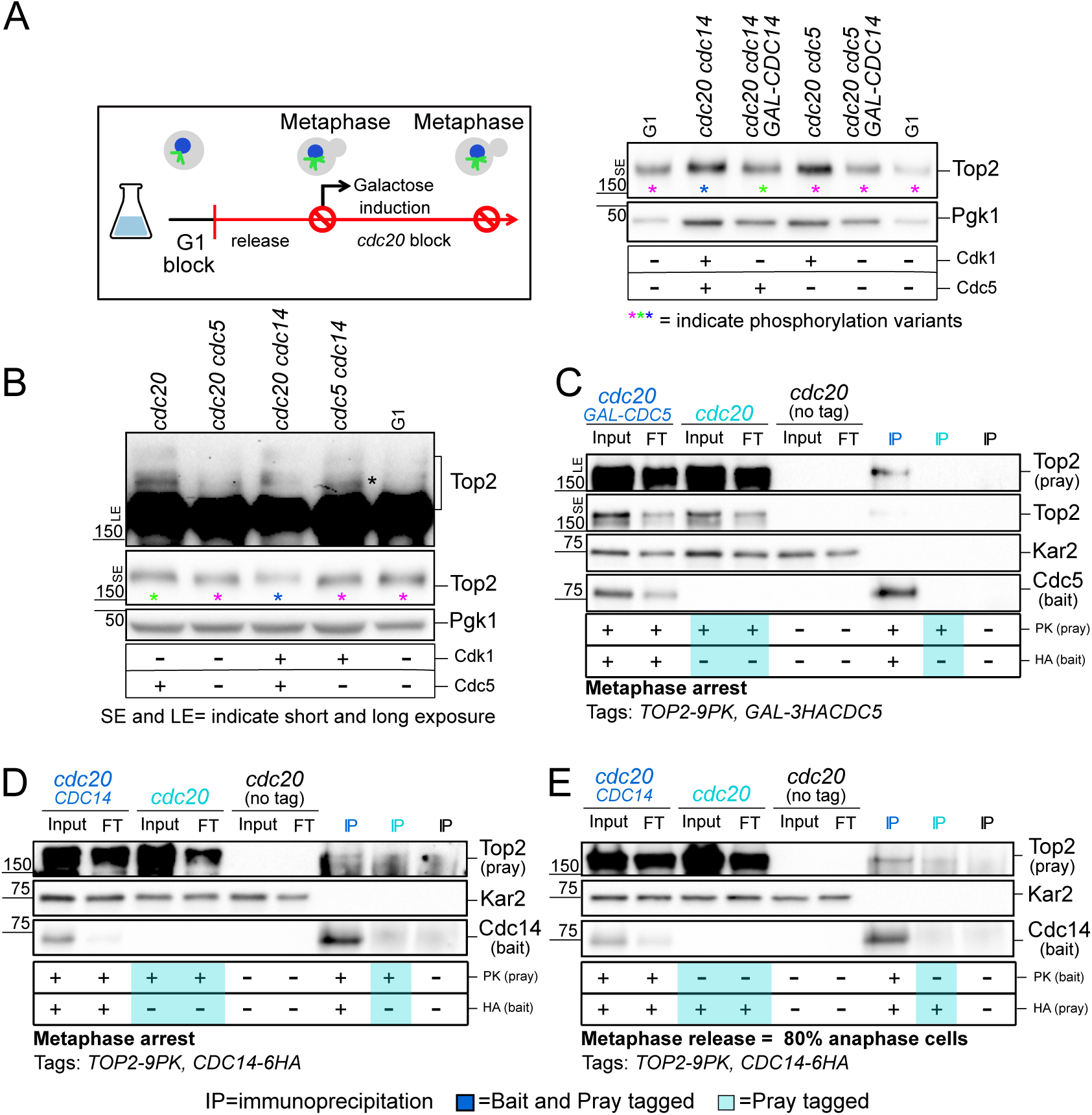
Cdc5 and Cdc14 affect Top2 phosphorylation. (**A**) *CDC20-AID cdc14-1* (Ry8782), *CDC20-AID cdc14-1 GAL-CDC14* (Ry11307), *CDC20-AID cdc5-as1* (Ry8785), *CDC20-AID cdc5-as1 GAL-CDC14* (Ry11241) were synchronously released from a G1 block into metaphase (*cdc20* arrest) in YPR medium. Upon arrest, galactose was added to induce Cdc14 expression, after which samples were collected samples were collected for immunostaining against Top2, and Pgk1 on SDS-PAGE. (**B**) *CDC20-AID* (Ry8315), *CDC20-AID cdc5-as1* (Ry8785), *CDC20-AID cdc14-1* (Ry8782) and *cdc5-as1 cdc14-1* (Ry7998) cells carrying a *TOP2-9PK* fusion were synchronously released from a G1 block in conditions restrictive for the proteins of interest. 180 minutes after release, samples were collected for immunostaining against Top2, and Pgk1 on SDS-PAGE. (**C**) *CDC20-AID* (Ry4852), *CDC20-AID TOP2-9PK* (Ry8315) and *CDC20-AID GAL-3HACDC5 TOP2-9PK* (Ry11294) cells were synchronously released from a G1 block into metaphase (*cdc20* arrest) in YPR medium. Upon arrest, galactose was added to induce Cdc5 expression, after which samples were collected for Cdc5-HA immunoprecipitation and analysed for interaction with Top2. Kar2 serves as loading control. (**D-E**) *CDC20-AID* (Ry4852), *CDC20-AID TOP2-9PK* (Ry8315) and *CDC20-AID CDC14-6HA TOP2-9PK* (Ry11279) were cells were released from a G1 block into metaphase (*cdc20* arrest) in YPR medium. At the arrest (**D**) or 45 minutes after release from the metaphase block, when 80% cells are in early anaphase (**E**), samples were collected for Cdc14-Ha immunoprecipitation and analysed for interaction with Top2. In the mutants of interest, the table reporting “+/- Cdk1 and Cdc5" indicates the presence or absence of phosphorylation events mediated by Cdk1 and/or Cdc5. Input, FT, flowthrough; IP, immunoprecipitate; IP+λPP, immunoprecipitate treated with λ phosphatase (PP).

To investigate the relative contribution of Cdc5 and Cdc14 on Top2 phosphorylation, we next looked at the effect of inactivating them separately in metaphase and in combination in mini-anaphase. In metaphase-arrested *cdc20* cells, bulk Top2 migrated more slowly than in G1 (Fig. 8B and Supplementary Fig. 11B, green *vs* magenta asterisk).

Loss of Cdc5 produced a faster, G1-like band, indicating that most mitotic phosphorylation depends on Cdc5 (Fig. 8B and Supplementary Fig. 11B). The same band was also detected in the *cdc5 cdc14* mutant, further supporting that Cdc5 plays a major role in regulating Top2 during metaphase. Longer exposures revealed additional, less abundant slower-migrating Top2 species in *cdc20* cells but absent in G1 and *cdc20 cdc5* mutants, indicating that Cdc5 also contributes to the additional modification of a specific subset of Top2 during metaphase. Notably, *cdc5 cdc14* cells exhibited a unique Top2 band (black asterisk) absent in *cdc20 cdc5* cells, suggesting that upon anaphase onset, Cdc14 dephosphorylates a pool of Top2 phosphorylated by a kinase other than Cdc5. The fact that inactivating Cdc14 during metaphase did not noticeably increase Top2 phosphorylation—bands appeared only slightly more diffuse, if at all (Fig. 8B and Supplementary Fig. 11B, blue asterisk, compare *cdc20* and *cdc20 cdc14*)—suggests that Cdc14 activity is largely restricted to anaphase. Since Cdc14 primarily reverses Cdk-dependent phosphorylation, these results point to Cdk as the kinase responsible for this Top2 modification. Consistent with a temporal specificity —Cdc5 acting primarily in metaphase and Cdc14 in anaphase—co-immunoprecipitation in *cdc20*-arrested cells revealed a strong Top2–Cdc5 interaction (Fig. 8C; Supplementary Fig. 11C), which was further enhanced using a kinase-dead substrate-trap allele (Supplementary Fig. 11D). In contrast, Cdc14 associated only weakly with Top2 during metaphase (Fig. 8D, Supplementary Fig. 11E) but showed a stronger interaction in early anaphase (Fig. 8E), consistent with a subsequent role in Top2 dephosphorylation.

Phosphoproteomic analyses of metaphase (*cdc20*), mini-anaphase (*cdc5 cdc14*), and anaphase (*cdc15*) cells (61) identified four Top2 residues—S1252, S1254, S1310, and T1314—whose phosphorylation levels varied across these stages, indicating dynamic *in vivo* regulation of Top2 during mitosis (Supplementary Fig. 11F). Among these, S1252 and S1254 lie within motifs compatible with phosphorylation by Cdk1 and Cdc5, respectively, with S1252 previously validated as a *bona fide* Cdk1 target (75–79). Their phosphorylation kinetics further support this interpretation, with S1252 behaving as a likely Cdc14 dephosphorylation target and S1254 as a Cdc5 phosphorylation site during metaphase. The latter is identified with higher confidence, consistent with the stronger effect of Cdc5 observed at this stage. Both residues map to the C-terminal domain (CTD) of Top2 - a regulatory hub that modulates enzyme function (76, 79) - and lie in close proximity to the SUMOylated lysines K1246 and K1247, which influence Top2 localization (73) (Supplementary Fig. 11G). Consistent with its role as a flexible signaling platform, the CTD is predicted to be intrinsically unstructured by AlphaFold3 (Supplementary Fig. 11G). Together, these findings reveal that Cdc5 and Cdc14 fine-tune Top2 phosphorylation through distinct yet coordinated mechanisms, integrating phospho- and SUMO-dependent control within the CTD to ensure proper Top2 activity during mitosis.

## Discussion

Genome integrity requires the complete removal of all physical linkages between sister chromatids during mitosis. While regulation of cohesin cleavage has been extensively characterized, the mechanisms that govern the resolution of DNA-based linkages remain less well understood. Using *cdc5 cdc14* budding yeast mutants, which arrest in mini-anaphase with a short spindle and unsegregated nuclei, we show that these cells accumulate sister chromatid intertwines (SCIs), primarily in the form of catenanes. Both Cdc5 and Cdc14 are required for their resolution, with Cdc5 playing a predominant role early in mitosis and Cdc14 acting more strongly during anaphase and specifically in rDNA segregation. This difference may explan why spindle elongation is more severely impaired in *cdc5* mutants compared to *cdc14* mutants (29).These two proteins exert their functions primarily through the regulation of Top2, the enzyme responsible for resolving DNA entanglements. Our findings indicate that Cdc5 and Cdc14 influence Top2 localization and modulate its post-translational modifications.

### Cdc5 and Cdc14 modify Top2 phosphorylation and SUMOylation

Our findings, together with previous work, confirm that Cdc5 promotes Top2 SUMOylation (32, 33). Although SUMOylation has been proposed to facilitate Top2 recruitment to chromatin (32, 33), we find that loss of Top2 SUMOylation—either specifically or as part of a global reduction in mitotic SUMOylation—only modestly affects the resolution of sister chromatid intertwines (SCIs). This suggests that SUMOylation may act in a locus- or context-specific manner rather than serving as a general requirement for decatenation.

Beyond SUMOylation, we show that Cdc5 and Cdc14 independently modulate Top2 phosphorylation. While our data establish a clear regulatory link, future mutagenesis studies will be necessary to pinpoint the specific residues involved. Cdc5 promotes, whereas Cdc14 opposes, Top2 phosphorylation, defining an antagonistic regulatory axis that ensures the precise tuning of Top2 activity required for faithful chromosome segregation. Importantly, our data also suggest that these two enzymes act on distinct Top2 pools: Cdc5 broadly modulates Top2 DNA binding and chromatin association, whereas Cdc14 functions more selectively, targeting specific Top2 subpopulations, such as those associated with the nucleolus. This spatial division of labor aligns with their partially redundant yet complementary roles in coordinating spindle elongation and chromosome segregation (29) and with the coexistence of phosphorylation and dephosphorylation events that fine-tune mitotic progression (80). The ability of Cdc5 to phosphorylate Top2 appears conserved across species, as human Plk1 similarly phosphorylates TOP2A *in vitro* (81).

The presence of key post-translational modifications within the Top2 C-terminal domain (CTD) underscores their critical role in regulating enzyme function. This may also explain why yeast Top2 proved less effective than its *Chlorella* counterpart, which lacks this domain.

Notably, in human TOP2A, the major SUMO acceptor lysine 1240 lies adjacent to phosphosites (T1244, S1247) whose modification modulates SUMO attachment (82), hinting at a conserved phospho-SUMO crosstalk mechanism that could control Top2 localization and activity. Although this form of regulation does not seem to account for the majority of Top2 regulation in yeast, we cannot exclude that it functions at specific sites.

Overall, our results establish Cdc5 and Cdc14 as regulators of Top2 post-translational modification. By tuning phosphorylation and SUMOylation dynamics, these enzymes coordinate Top2’s chromatin engagement and decatenation function, providing a mechanistic link between mitotic regulation and the maintenance of genome integrity.

### A model for Cdc5 and Cdc14 mediated regulation of Top2 localization

Top2 is essential for resolving catenanes throughout the cell cycle. In S phase, it tracks replication forks, supporting fork progression by limiting supercoil accumulation, which converts to catenanes at replication termination (83, 84). Its role in catenane resolution extends into metaphase, guided by chromosome condensation and spindle attachment, and culminates in anaphase, when cohesin cleavage allows full decatenation. In late anaphase, Top2 removes residual catenanes that would otherwise persist as anaphase bridges (4). Our data show that loss of Cdc5 and Cdc14 from metaphase onward blocks SCI resolution, indicating that their function is critical in metaphase and anaphase rather than in S phase.

We have shown that, in metaphase, before spindle attachment, Top2 is enriched at centromeres, consistent with yeast and mammalian studies (15) (85). We found that tension dramatically reduces this centromeric enrichment, coinciding with reported conformational rearrangements of pericentromeric chromatin (38) and the delocalization of cohesin, shugoshin, condensin, PP2A-Rts1, and the Chromosomal Passenger Complex (15, 86, 87). Unlike condensin, whose levels rise along chromosome arms once tension is established, Top2 did not show a similar redistribution in our assays, suggesting that its recruitment is temporally distinct and not strictly coupled to condensin. Future investigations should elucidate the interactions between these factors and Top2, illuminating the mechanisms that govern its recruitment and subsequent delocalization.

How Top2 is guided in anaphase is less clear. In budding yeast, while it is known that catenanes persist until cohesin cleavage, the collaboration between Top2 and condensin in resolving these structures remains largely unexplored. Top2 recruitment has so far been described mainly on ultrafine anaphase bridges, where Dpb11/TOPBP1 directs its localization (66, 88). Our ChIP-seq showed that Top2 levels are reduced in *cdc5 cdc14* mini-anaphase arrest compared to metaphase, both at centromeres and along chromosome arms, consistent with the defects in chromatin recruitment and nuclear localization observed in the double mutant. This points to an essential role for Cdc5 and Cdc14 in sustaining Top2 function. Wild-type cells induced to segregate chromosomes by artificial cohesin cleavage resolve catenanes efficiently, indicating that the machinery for resolution is already present in metaphase, reinforcing the idea that Cdc5 and Cdc14 act at this stage. However, their contribution continues into anaphase, as shown by the persistence of SCIs in single mutants even with fully elongated spindles.

Because most Cdc14 remains sequestered in the nucleolus during metaphase, Cdc5 likely provides the primary input at this stage, promoting phosphorylation and SUMOylation events that recruit Top2 to centromeres. This is supported by our co-immunoprecipitation data showing a strong Cdc5–Top2 interaction in metaphase that is further enhanced in a kinase-dead substrate-trap allele, while Cdc14 associates weakly at this stage but binds more strongly in early anaphase. These findings suggest that Cdc5 initiates Top2 activation in metaphase, whereas Cdc14 fine-tunes its activity during anaphase by dephosphorylating specific residues. Acting together, Cdc5 and Cdc14 coordinate Top2 function along chromatids, coupling it to condensin-driven chromosome recoiling. They may also influence Dpb11–Top2 interactions, reminiscent of human TOPBP1–Top2A regulation, where the BRCT domain of TOPBP1 recognizes phosphorylated substrates—raising the possibility that Cdc5 phosphorylation facilitates Top2 recruitment to anaphase bridges (66, 88).

### A comprehensive model for sister chromatid separation

Top2-mediated catenane resolution requires condensin which compacts chromatids and exposes entanglement sites. We confirm that condensin is essential for Top2 function and that the catenane accumulation in *cdc5 cdc14* cells is not due to condensin inactivity. Although Cdc5 and Cdc14 also promote spindle elongation (29), forcing elongation in *cdc5 cdc14* does not bypass SCI defects, and anaphase bridges persist in *cdc5* and *cdc14* single mutants with fully extended spindles. Thus, their contribution extends beyond spindle mechanics likely to direct regulation of Top2 localization and activity.

Beyond catenanes, Cdc5 and Cdc14 contribute to the resolution of replication and recombination intermediates. Cdc5 activates the nuclease Mms4-Mus81 in metaphase and Cdc14 the nuclease Yen1 in anaphase (4). Mms4-Mus81 and Yen1 are backup mechanism that resolve replication and recombination intermediates that escape S phase or originate from the anaphase replication of difficult to replicate regions (63). In addition, Cdc5 and Cdc14 facilitate cohesin cleavage: Cdc5 regulates the anaphase-promoting complex (APC/C) to ensure timely degradation of the cohesin subunit Scc1, while Cdc14 enhances securin degradation by counteracting its phosphorylation (89).

Together, these functions eliminate all classes of chromatid linkages while coordinating their removal with spindle elongation, which acts as a “molecular ruler” that couple enzymatic resolution with mechanical separation (Fig. 9)

**Fig. 9:**
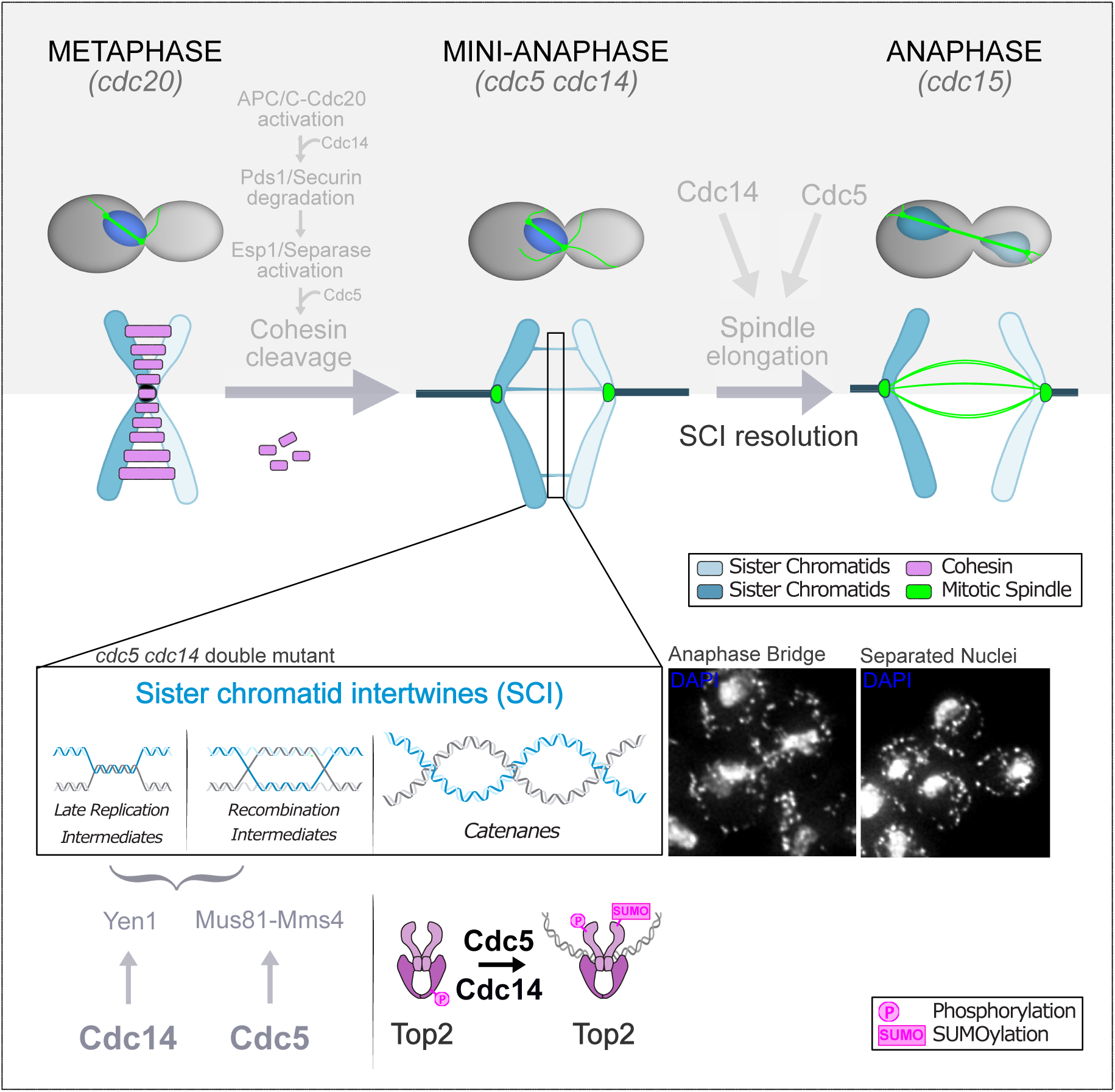
Schematic model highlighting the central roles of Cdc14 and Cdc5 in driving sister chromatid separation and genome stability during mitosis. Cdc5 and Cdc14 orchestrate chromosome segregation through multiple mechanisms. They regulate protein-mediated linkages, such as cohesin cleavage via control of APC/C and securin degradation, and DNA-mediated sister chromatid intertwines (SCIs). Central to the latter, they promote Top2-dependent decatenation—by modulating its post-translational modifications and localization—acting in parallel to condensin activity. Additionally, they facilitate the resolution of replication and recombination intermediates through activation of nucleases Mms4-Mus81 (Cdc5) and Yen1 (Cdc14). By coordinating the resolution of diverse chromatid linkages with active promotion of spindle elongation, they ensure proper segregation and genomic stability during mitosis.

### DNA Intertwines and Mitosis: Friends or Foe?

SCIs are unavoidable byproducts of DNA replication, but their persistence into mitosis (4), raises the possibility of a physiological role in cohesion or segregation timing. While SCIs can compensate for cohesin loss in some contexts (10) (90), whether this occurs during an unperturbed cell cycle remains unclear. It is tempting to speculate that transient SCIs provide resistance to spindle forces, delaying separation until segregation is fully coordinated. If left unresolved, however, SCIs become anaphase bridges that threaten genome stability. Spindle forces rarely break chromosomes, but cytokinesis can sever chromatin bridges in both yeast and mammals (19, 91, 92). In mammalian cells, this can result in fragmentation or tetraploidy, and cells rely on the abscission checkpoint—or NoCut pathway in yeast — to delay cytokinesis in the presence of bridges (93, 94). Strikingly, this checkpoint uses Topoisomerase II localization to chromatin as a sensor of DNA bridges, coupling separation with abscission (95). Our data reveal a complementary safeguard: a Cdc5–Cdc14–Top2 module that couples SCI resolution to spindle elongation, ensuring that chromatids are disentangled before cell division completes and thereby preserving genome stability.

## Supporting information

Supplementary informations

## Data Availability

Data are available in the main text or the supplementary materials. The Chip-seq data have been deposited to the GEO repository (GSE294907). The following secure token has been created to allow review of record GSE294907 while it remains in private status: sdebyikwhzqnvqn

## Funding

The work of the Visintin’s laboratory was supported by the Italian Association for Cancer Research, AIRC (IG-2012-12878 and IG-2015-16886); the Italian Ministry of Health (RF-2011-02347470) and in part by an International Early Career Scientist grant from the Howard Hughes Medical Institute. AF was supported by the FIRC-AIRC “Luigi, Antonietta e Gabriele Gerosa” fellowship. LFM was supported by a fellowship of FIRC “Titino Colombo”. AD was supported by the FIRC-AIRC “Luca Erizzo” fellowship. AF, AD and LFM were PhD students within the European School of Molecular Medicine (SEMM). This work was partially supported by the Italian Ministry of Health with Ricerca Corrente and 5×1000 funds.

## Conflict of interest

The authors declare no competing interests.

## Acknowledgements

We thank Dana Branzei, Frank Uhlmann, Steve West, Stephen Elledge, Xiaoxue Snow Zhou, Chiara Tordonato, Simone Tamburri and Tomo Tanaka for strains and reagents; Frank Uhlmann and members of the Visintin laboratory for helpful discussions; Alessandro Vetere for his help with the analysis of IF data, Adèle Marston for help with the Chip-seq experiments and Daniel Fernández-Pérez for support in the analysis of the experiment; Federico Zucca for the phospho-proteomic analysis; Giuseppe Ciossani of the Biochemistry and Structural Biology Unit (BSU) and Simona Rodighiero of the Imaging Unit at the IRCCS European Institute of Oncology (IEO, Milan, Italy) for structure prediction and depiction and image analysis assistance, respectively.

## Author contributions

AF, LFM, and RV conceived and designed the study. AF and LFM performed most of the original experimental work, with support from CV and EC. For the revised version, CV, AD, and EC conducted the new experiments and data analyses. LFM carried out the bioinformatics analyses. AF, LFM, and RV wrote the manuscript and prepared the figures, with input from EC, AD, and CV. All authors read and approved the final manuscript.

## Notes

### Competing Interest Statement

The authors have declared no competing interest.

