## Supplementary informations for "Safeguarding Genome Integrity: Polo-like kinase Cdc5 and phosphatase Cdc14 Orchestrate Topoisomerase II-Mediated Catenane Resolution in Mitosis": Massari_Finardi_Supplementary copy.pdf

**This PDF file includes eleven supplementary figures and four tables:**

Supplementary Figures 1 to 11

Supplementary Table 1: Yeast strains used in this study
Supplementary Table 2: Centromere coordinates
Supplementary Table 3: Mean nuclear Top2 fluorescence intensity
Supplementary Table 4: Coefficient of variation

A

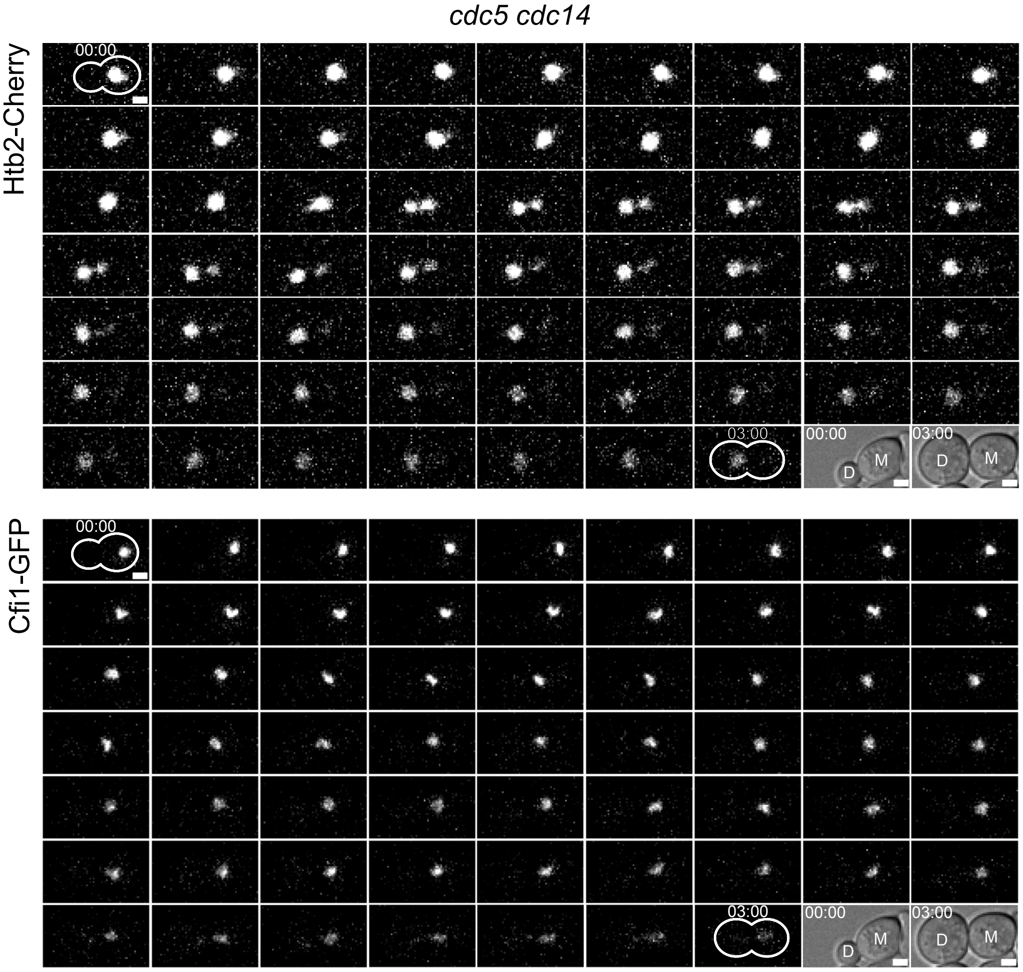


B

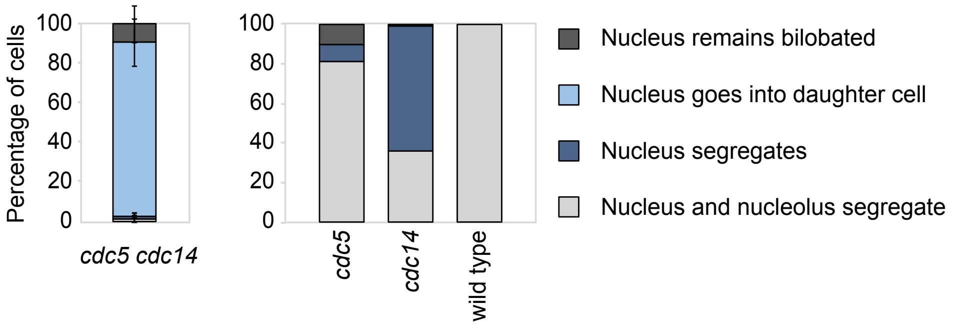


Supplementary Figure 1: ***cdc5 cdc14* nuclei move as a whole into daughter cells.** (**A**-**B**) *cdc5-as1 cdc14-1* (Ry6099), *cdc5-as1* (Ry6289), *cdc14-1* (Ry6101) and wild-type (Ry6105) cells, all carrying *HTB2-Cherry CFI1-GFP* fusions were synchronously released from a G1 block in conditions restrictive for *cdc5* and *cdc14*. Cells were imaged starting from metaphase every 3 minutes for 3 hours. Nuclear and nucleolar morphologies were scored. Scale bars = 2 µm (**A**). Cells were assigned to the indicated categories according to nuclear and nucleolar segregation during the time-lapse (**B**).

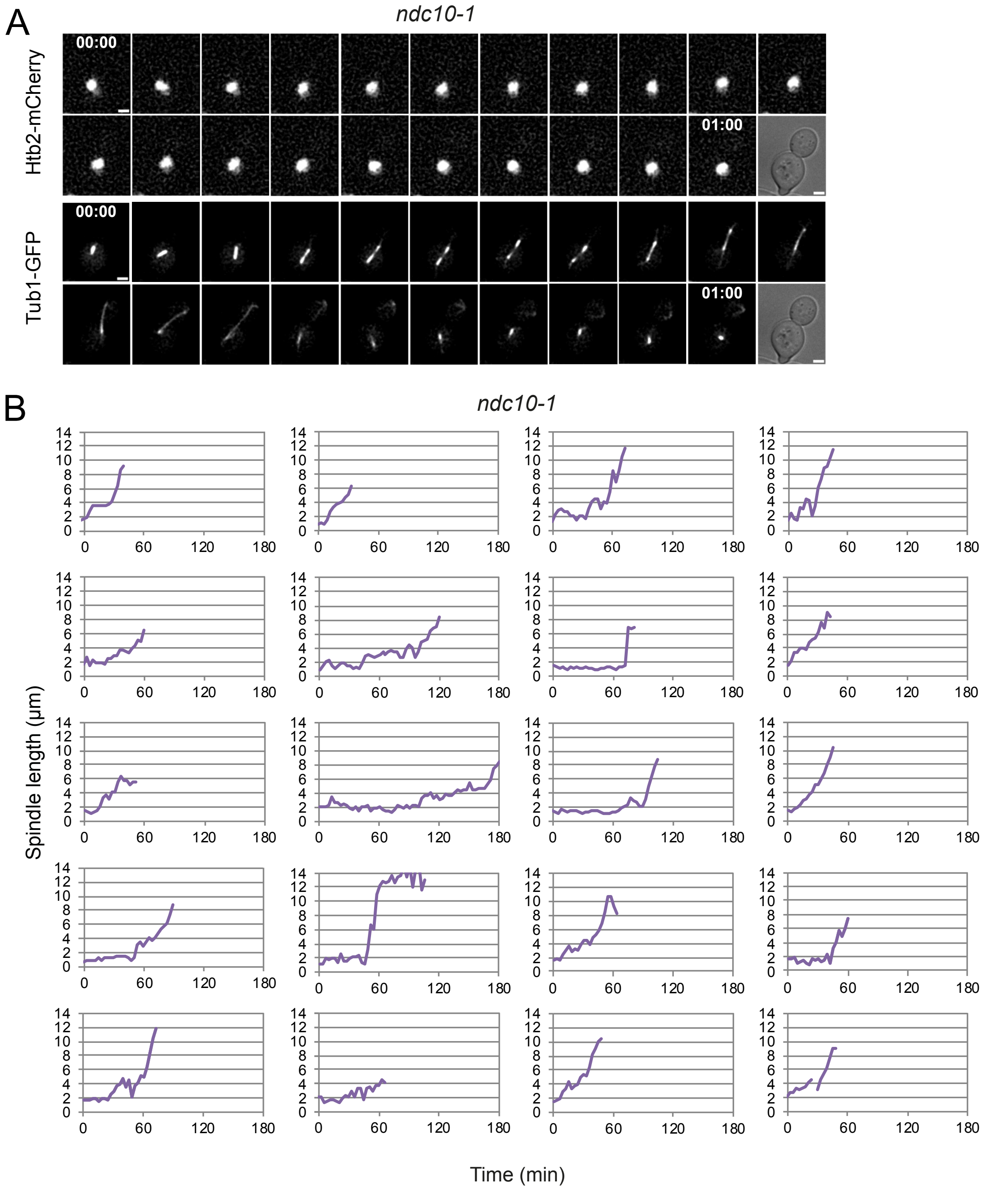


Supplementary Figure 2: ***ndc10* cells uncouple chromosome segregation and spindle elongation.**
*ndc10-1* (Ry6590) cells carrying a *HTB2-Cherry* and a *GFP-TUB1* fusion were synchronously released from a G1 block at 37°C (restrictive for *ndc10*). Cells were imaged starting from metaphase every 3 min for 3 hours. Representative images of the first hour of imaging are shown. Nuclear and spindle morphologies were scored. Scale bars = 2 µm (**A**). Spindle length was measured throughout the timelapse on x,y,z (**B**). n=20 cells

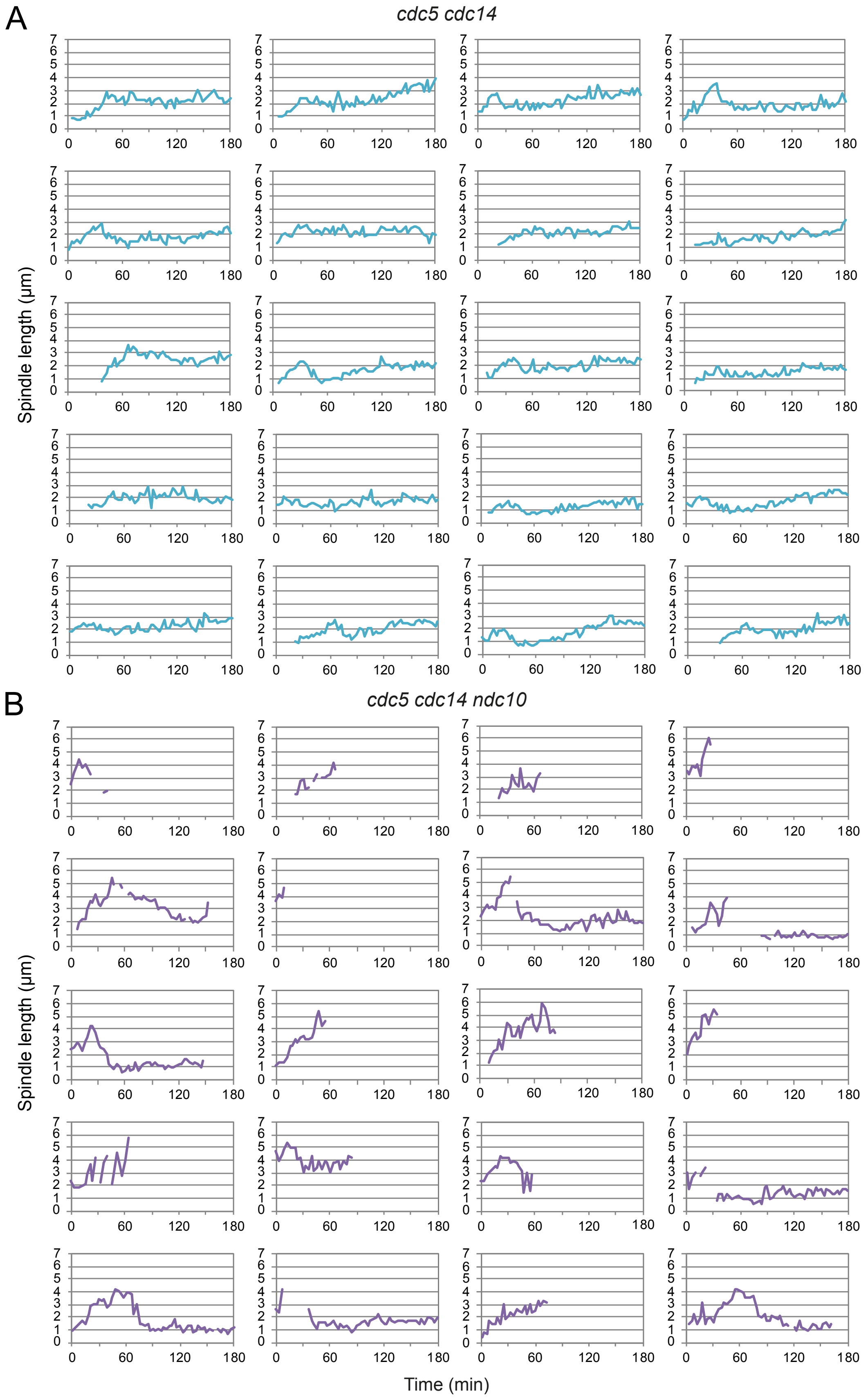
A
cdc5 cdc14
B
cdc5 cdc14 ndc10
Spindle length (µm)
Time (min)
0
60
120
180
0
1
2
3
4
5
6
7

Supplementary Figure 3: ***cdc5 cdc14* cells retain residual cohesion between sister chromatids.**

*cdc5-as1 cdc14-1* (Ry6591- **A**) and *cdc5-as1 cdc14-1 ndc10-1* (Ry6589 - **B**) cells carrying a *HTB2-Cherry* and a *GFP-TUB1* fusion were synchronously released from a G1 block in conditions restrictive for *cdc5*, *cdc14*, and *ndc10*. Cells were imaged starting from metaphase every 3 min for 3 hours. n=20 cells for each strain. Spindle length was measured throughout the timelapse on x,y,z.

Massari and Finardi_Supplementary Figure 4

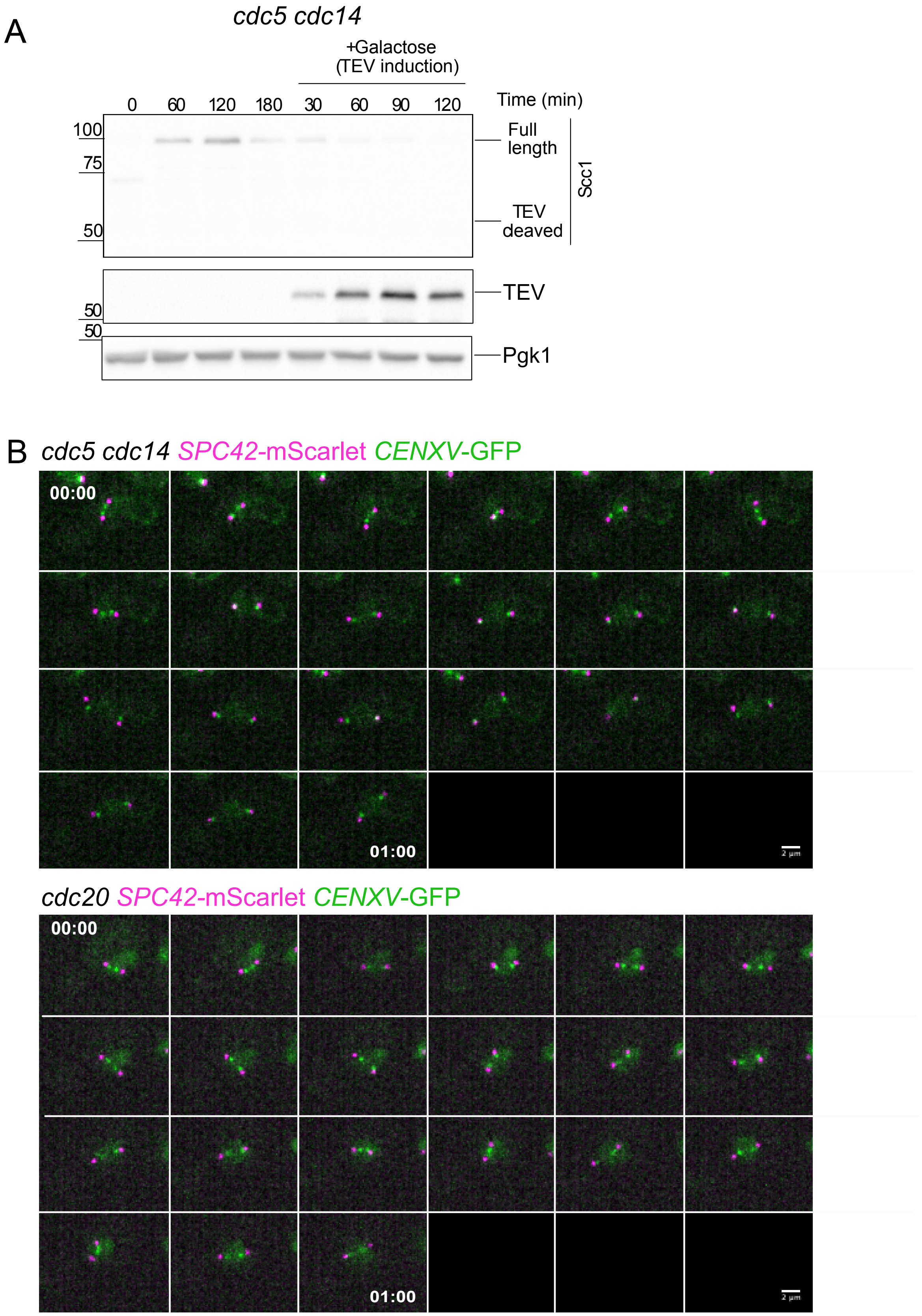

Supplementary Figure 4: **Cohesin is lost in *cdc5 cdc14* cells**
(**A**) *cdc5-as1 cdc14-1 SCC1-TEV GAL-TEV* (Ry2795) cells were synchronously released from a G1 block in YPR in conditions restrictive for the mutant alleles. At the arrest (3h after release), galactose was added to induce TEV protease expression. Samples were collected at the indicated time after induction and analyzed by Western blotting to assess protease expression (TEV) and cohesin cleavage, probing the Scc1 subunit of the cohesion complex. Pgk1 was used as loading control.
(**B**) *cdc5-as1 cdc14-1* (Ry11247) and *CDC20*-AID (Ry11273) cells carrying *SPC42-mScarlet* and *CENXV-GFP* fusions were synchronously released from a G1 block in conditions restrictive for the mutant alleles. Cells were imaged starting from metaphase every 3 minutes for 1 hours. Representative cells are shown. Scale bars = 2 μm.

Massari and Finardi_Supplementary Figure 5

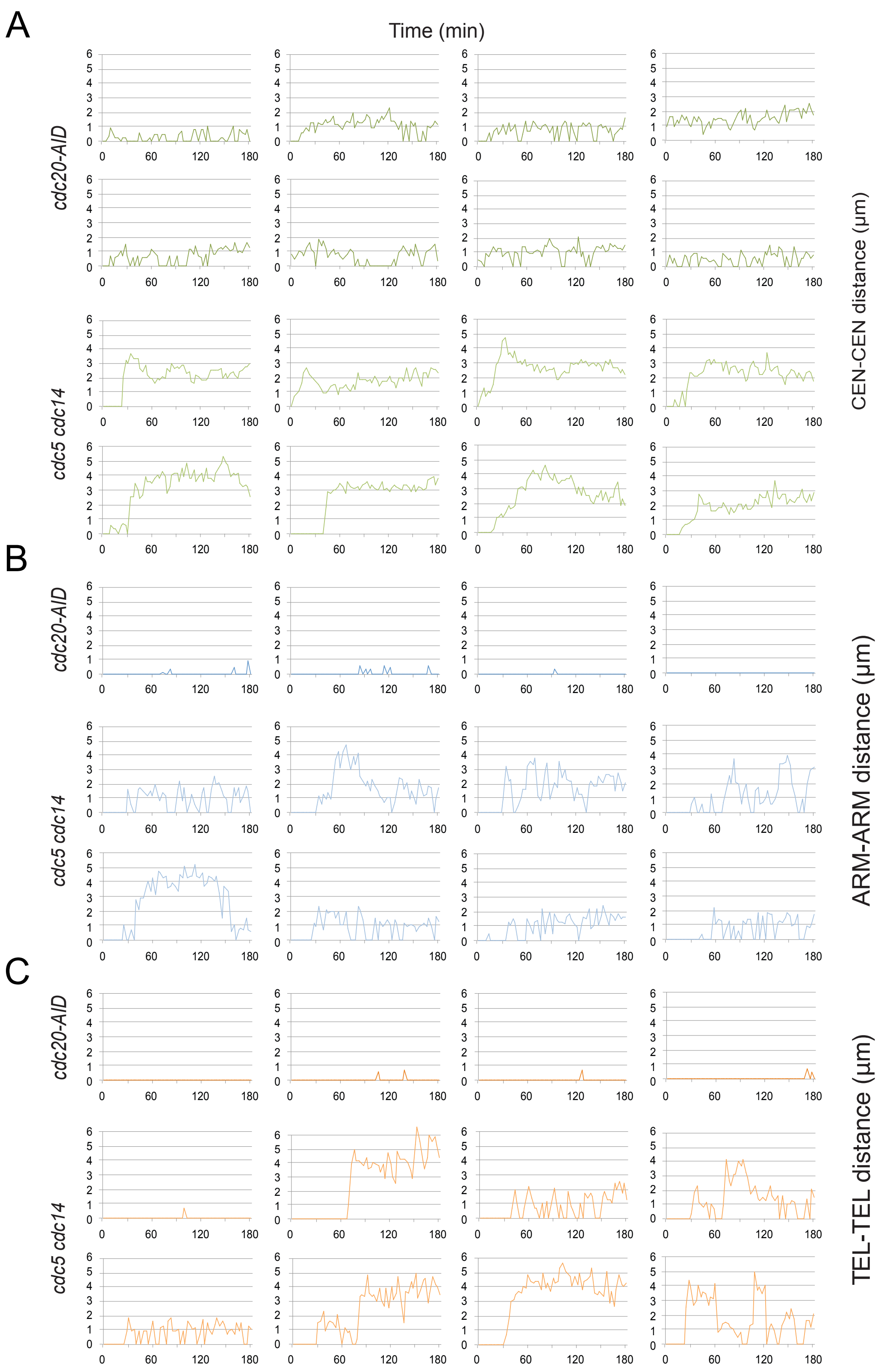

Supplementary Figure 5: ***cdc5 cdc14* retains some linkages at arms and telomeres.** *CDC20-AID* and *cdc5-as1 cdc14-1* cells carrying a: *CENXV-GFP (*Ry7519 and Ry5815*)* (**A**), *HIS3-GFP* (Ry7411 and Ry5824) (**B**) or *TELXV-GFP* (Ry7481 and Ry7098) (**C**) fusions were synchronously released from a G1 block into conditions restrictive for *cdc20*, *cdc5* and *cdc14*. Cells were imaged starting from metaphase every 3 minutes for 3 hours. Distance between sister GFP-dots was measured throughout the time-lapse on x,y,z. Representative cells are shown. n=20 cells were analyzed in each strain.

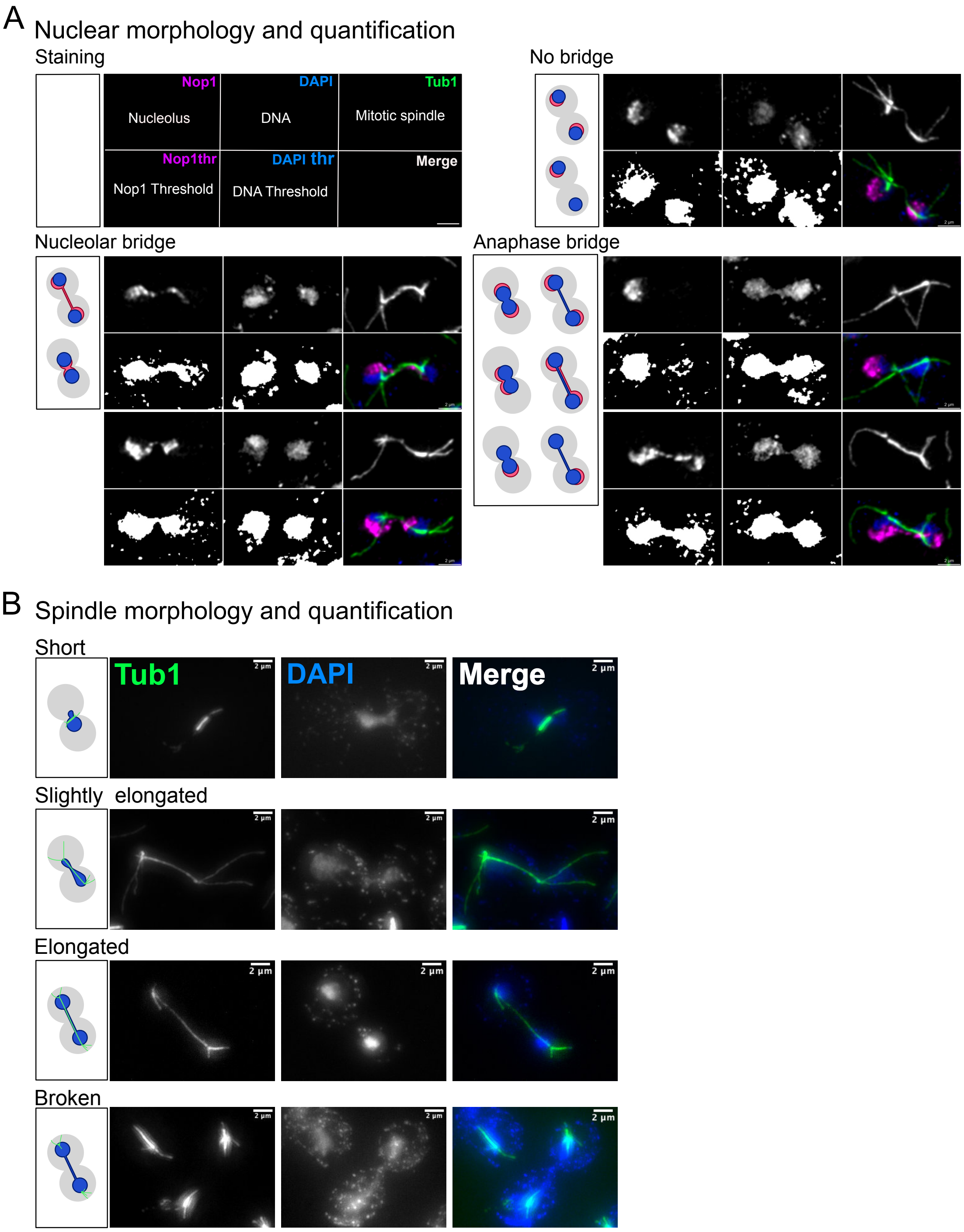


Supplementary Figure 6: **Nuclear and spindle morphologies.**
Representative cells from the experiment in Fig. 3 are displayed to illustrate the analyses. (**A**) Nuclear and nucleolar morphologies: a box indicates a schematic representation of the immunofluorescence (IF) images for nuclear and nucleolar categories. The top row includes, from left to right, raw grayscale images of Nop1 (nucleolus), DAPI (nucleus), and Tub1 (mitotic spindle). The second row shows, from left to right, the thresholded images used for the quantification of nucleolar (Nop1) and nuclear (DAPI) bridges. The final image combines

all channels, with Nop1 in magenta, DAPI in blue, and the mitotic spindle in green. Scale bars = 2 µm.
(**B**) Spindle morphology: a box indicates a schematic representation of the immunofluorescence (IF) images for spindle categories. For each category: from left to right, raw grayscale images of Tub1 (mitotic spindle) and DAPI (nucleus). The final image combines all channels, with the mitotic spindle in green and the DAPI in blue. Scale bars = 2 µm.

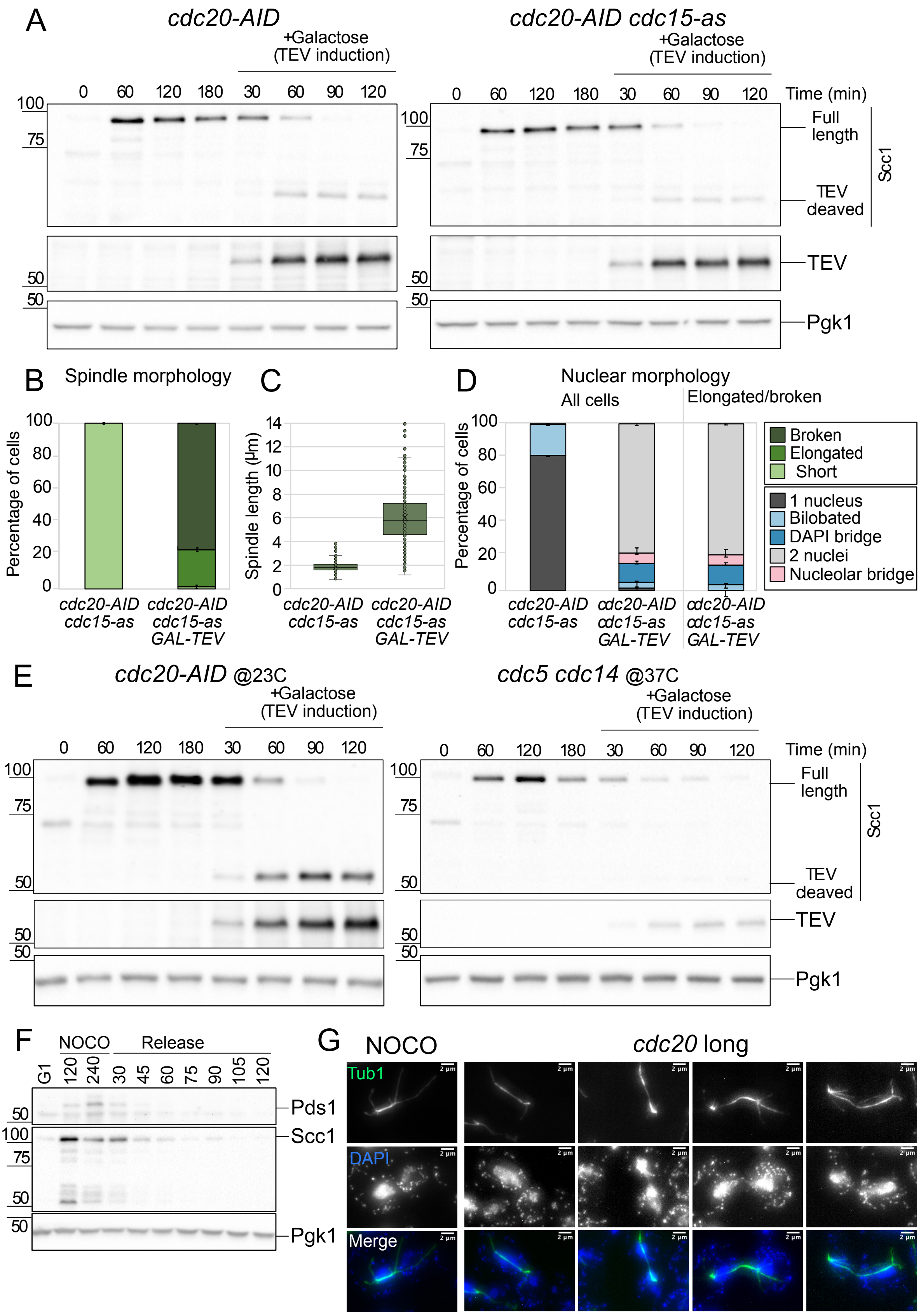
A
cdc20-AID
cdc20-AID cdc15-as
+Galactose (TEV induction)
0 60 120 180 30 60 90 120
Time (min)
100
75
50
Full length
TEV deaved
Scc1
TEV
Pgk1
B
Spindle morphology
Percentage of cells
cdc20-AID cdc15-as
cdc20-AID cdc15-as GAL-TEV
C
Spindle length (µm)
D
Nuclear morphology
All cells
Elongated/broken
Broken
Elongated
Short
1 nucleus
Bilobated
DAPI bridge
2 nuclei
Nucleolar bridge
E
cdc20-AID @23C
cdc5 cdc14 @37C
F
NOCO
Release
G1
Pds1
G
NOCO
cdc20 long
Tub1
DAPI
Merge
2 µm

Supplementary Figure 7: **Conditions for sister chromatid intertwine resolution are set in metaphase and require a functional bipolar spindle.**
(**A**-**D**) *CDC20-AID SCC1-TEV GAL-TEV* (Ry4931) and *CDC20-AID cdc15-as1 SCC1-TEV GAL-TEV* (Ry7563) cells were synchronously released from a G1 block in YPR in conditions restrictive for *cdc20*. At the metaphase arrest (3h after release), galactose was added to induce TEV protease expression. Samples were collected at the indicated time after induction and analyzed by western blotting to assess protease expression (TEV) and cohesin cleavage, probing the Scc1 subunit of the cohesion complex (**A**); and by IF (anti-Tub1, DAPI) to monitor spindle morphology (**B**), spindle lenght (**C**) and nuclear morphology (**D**). n=200 cells were analyzed for each condition. Pgk1 is used as loading control.
To compare Scc1 levels, samples from the *CDC20-AID SCC1-TEV GAL-TEV* strain were run side by side with samples from the *cdc5-as1 cdc14-1 SCC1-TEV GAL-TEV* strain (Supplementary Fig. 4) (**E**).
(**F-G**) *cdc5-as1 cdc14-1* (Ry1602) and *cdc5-as1 cdc14-1 MET-CDC20* (Ry3203) cells were synchronously released from a G1 block into a metaphase arrest by adding the depolymerizing drug, nocodazole, and methionine, respectively. When the arrest was complete, in nocodazole-treated cells Cdc5 and Cdc14 were inhibited for 45 minutes before release (NOCO). Instead, *MET-CDC20* cells were kept in the metaphase arrest for an additional 30 minutes (*cdc20* long) before inactivating the two proteins and next treated as above. A schematic of the rationale and experimental setup is shown in Fig. 3G
Samples were taken at the indicated times to probe Pds1 degradation and Scc1 cleavage to monitor SAC inactivation and anaphase entry (**F**). Representative images of the 120 minutes timepoint after release from Nocodazole or *MET-CDC20* cells kept longer in the metaphase block (*cdc20* long) are shown. The merge image shows the mitotic spindle in green and the DAPI in blue. Scale bars = 2 µm (**G**)

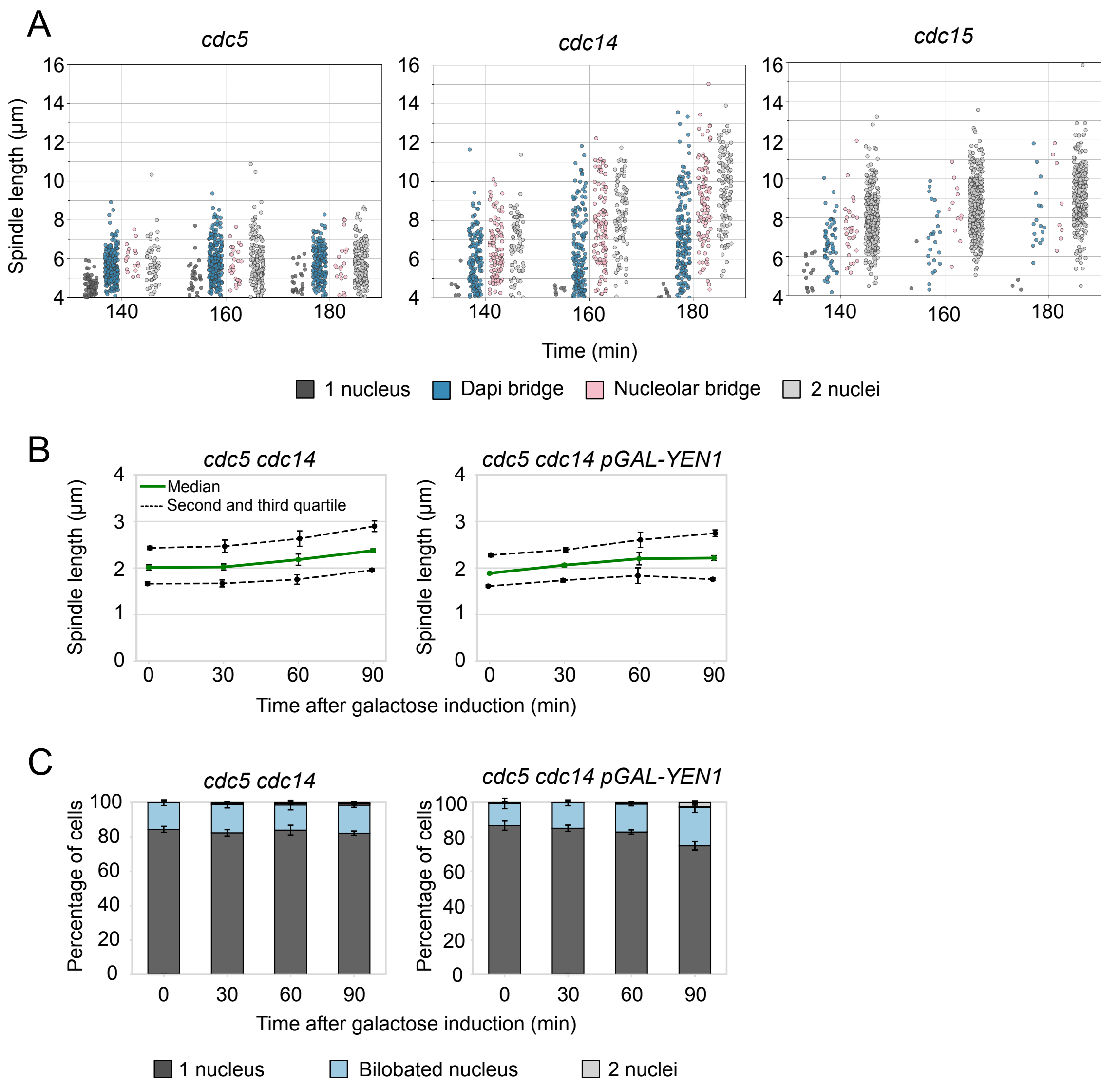


Supplementary Figure 8: **Yen1 overexpression does not rescue the *cdc5 cdc14* phenotype.**
(**A**) *cdc5-as1* (Ry2446), *cdc14-1* (Ry1573) and *cdc15-as1* (Ry1112) cells jitter plots for the experiment in Fig. 4 are shown. Each graph aggregates the cells from the three replicates of the experiment. Only cells with an anaphase spindle length (above 4 micrometers) are shown, consistent with the data presented in the histogram plots.
(**B-C**) *cdc5-as1 cdc14-1* (Ry1602) and *cdc5-as1 cdc14-1 pGAL-YEN1* (Ry10270) cells were synchronously released from a G1 block in YPR in conditions restrictive for *cdc5* and *cdc14*. At the *cdc5 cdc14* arrest (3h30 after release), galactose was added to induce Yen1 expression. Samples were collected at the indicated times after induction and analyzed through IF (anti-Tub1, DAPI) to monitor spindle length (**B**) and nuclear segregation (**C**). Three independent experiments are shown. n=100 cells were analyzed at each time point in each experiment. Spindle: Median (solid green line), second and third quartile (dotted lines) and s.e.m. (error bars) are shown (**B**). Nuclei: Mean and s.e.m. (error bars) are shown (**C**).

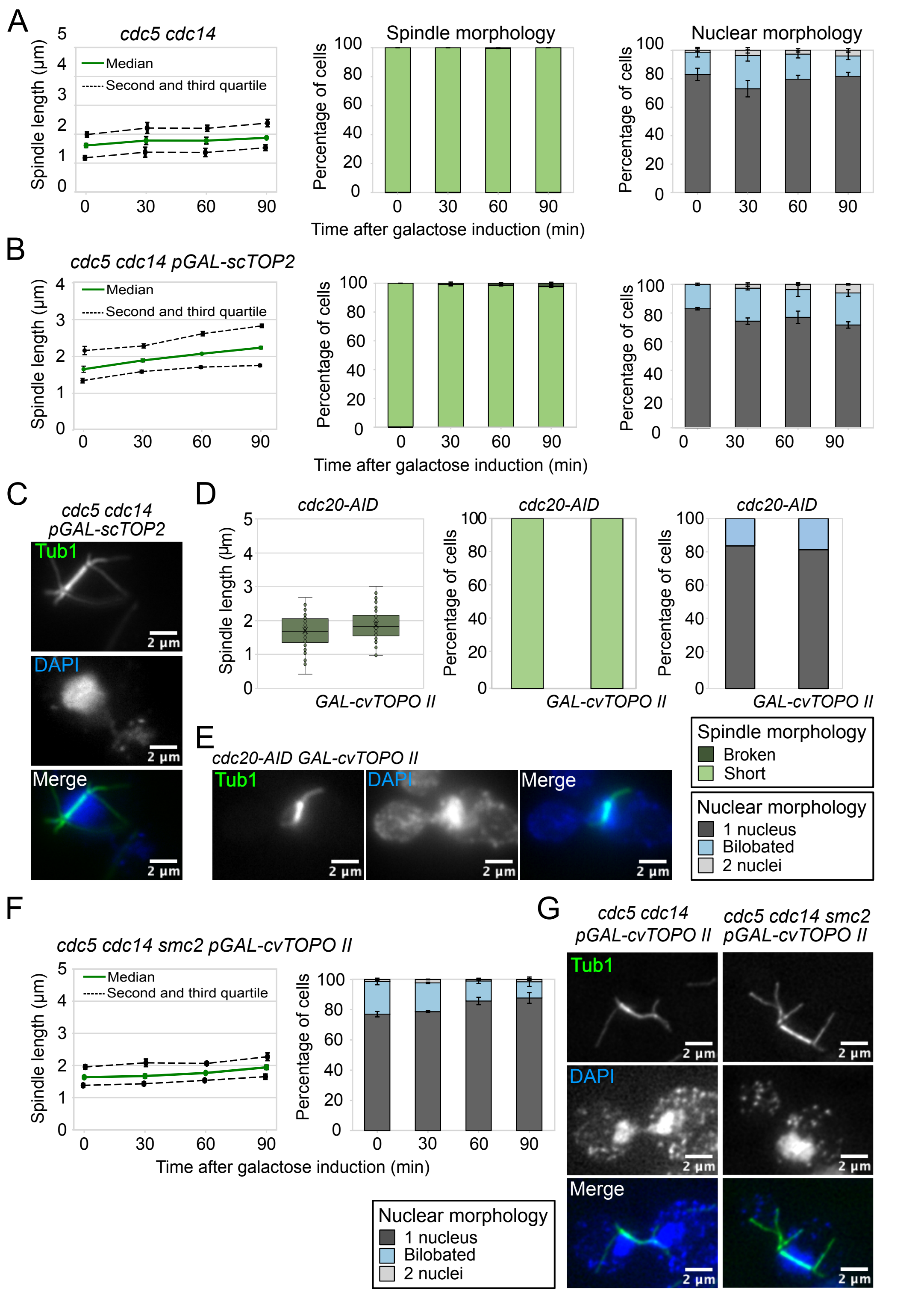


Supplementary Figure 9: **Condensin is active in *cdc5 cdc14* cells.**
*cdc5-as1 cdc14-1* (Ry1602 - **A**) and *cdc5-as1 cdc14-1 pGAL-scTOP2* (Ry5172 - **B**) cells were synchronously released from G1 block in YPR in conditions restrictive for *cdc5* and *cdc14.* At the terminal arrest (3h30 after release), galactose was added to induce Top2

overexpression. Samples were collected at the indicated time points after induction and analyzed through IF (anti-Tub1, DAPI) to monitor spindle length, morphology and nuclear morphology. Three independent experiments are shown. n=100 cells counted for each time point in each experiment. Median (solid line), second and third quartile (dotted lines) and s.e.m. (error bars) are shown. Mean and s.e.m. (error bars) are shown (**A-B**). Representative images are shown. Scale bars = 2 μm (**C**).
(**D-E**) *CDC20-AID* (Ry7874) and *CDC20-AID GAL-cvTOPOII* (Ry11225) cells were treated as above. Samples were collected 120 minutes after induction to monitor spindle length, morphology and nuclear morphology (**D**). Three independent experiments are shown. n=100 cells counted for each time point in each experiment. Mean and s.e.m. (error bars) are shown. Representative images in (**E**).
(**F-G**) *cdc5-as1 cdc14-1 smc2-8 GAL-cvTOPOII* (Ry7037) cells were synchronously released from G1 block in YPR in conditions restrictive for *cdc5*, *cdc14* and *smc2*. At the terminal arrest (3h30 after release), galactose was added to induce cv-TopoII overexpression. Samples were collected and probed for spindle length and nuclear morphology. Representative cells are shown. Three independent experiments are shown. n=100 cells counted at each time point in each experiment. Median (solid green line), second and third quartile (dotted lines) and s.e.m. (error bars) are shown. For the spindles. Mean and s.e.m. (error bars) are shown for nuclei (**F**). Representative images are shown. Scale bars = 2 μm (**G**).

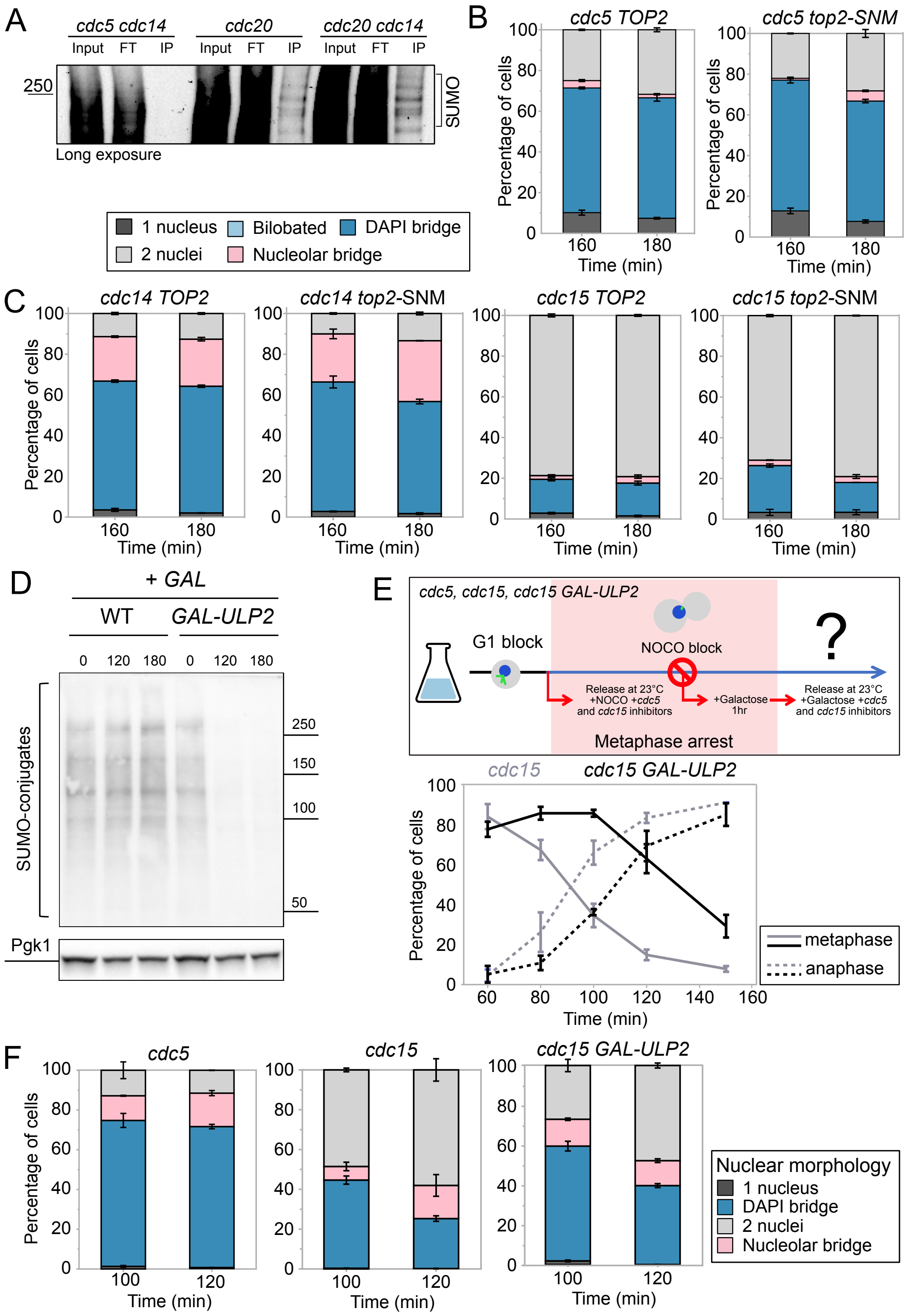


Supplementary Figure 10: **Lack of Top2 SUMOylation does not cause sister chromatid separation defects.**

(**A**) A long exposure of western blot in Fig. 7B is shown.

(**B-C**) *cdc5-as1 TOP2-3HA* (Ry10476), *cdc5-as1 top2-SNM-3HA* (Ry10482), *cdc14-1 TOP2-HA* (Ry10473), *cdc14-1 top2-SNM-HA* (Ry10485), *cdc15-as1 TOP2-HA* (Ry10539) and *cdc15-as1 top2-SMN-HA* (Ry10542) were synchronously released from G1 block in conditions restrictive for the proteins of interest. Samples were collected at the indicated times after release and analyzed through IF to score for the presence of anaphase bridges. At least 100 anaphase cells (spindle > 4 µm) were analyzed for each time point. Mean and s.e.m. (error bars) from three independent experiments are shown.
(**D**) Wild-type (Ry1) and *GAL-ULP2* (Ry8129) cells were grown in YPR to exponential phase. Galactose was added to induce Ulp2 overexpression. Samples were collected at the indicated times after induction for western blot. Total protein extracts were probed with an anti-SUMO antibody.
(**E**-**F**) *cdc5-as1* (Ry2446), *cdc15-as1* (Ry1112) and *cdc15-as1 GAL-ULP2* (Ry10532) cells were synchronously released from G1 block in YPR in conditions restrictive for *cdc5* and *cdc15* and in presence of nocodazole. At the metaphase arrest (3h post-release), galactose was added to induce Ulp2 expression. 1h after induction, cells were released from the meta block into the *cdc5* and *cdc15* arrest, in fresh medium without nocodazole in presence of high levels of Ulp2. Samples were collected at the indicated times after release from metaphase and analyzed through IF to monitor cell cycle progression through spindle (**E**), nuclear and nucleolar morphologies (**F**). n=100 cells counted at each time point in each experiment.

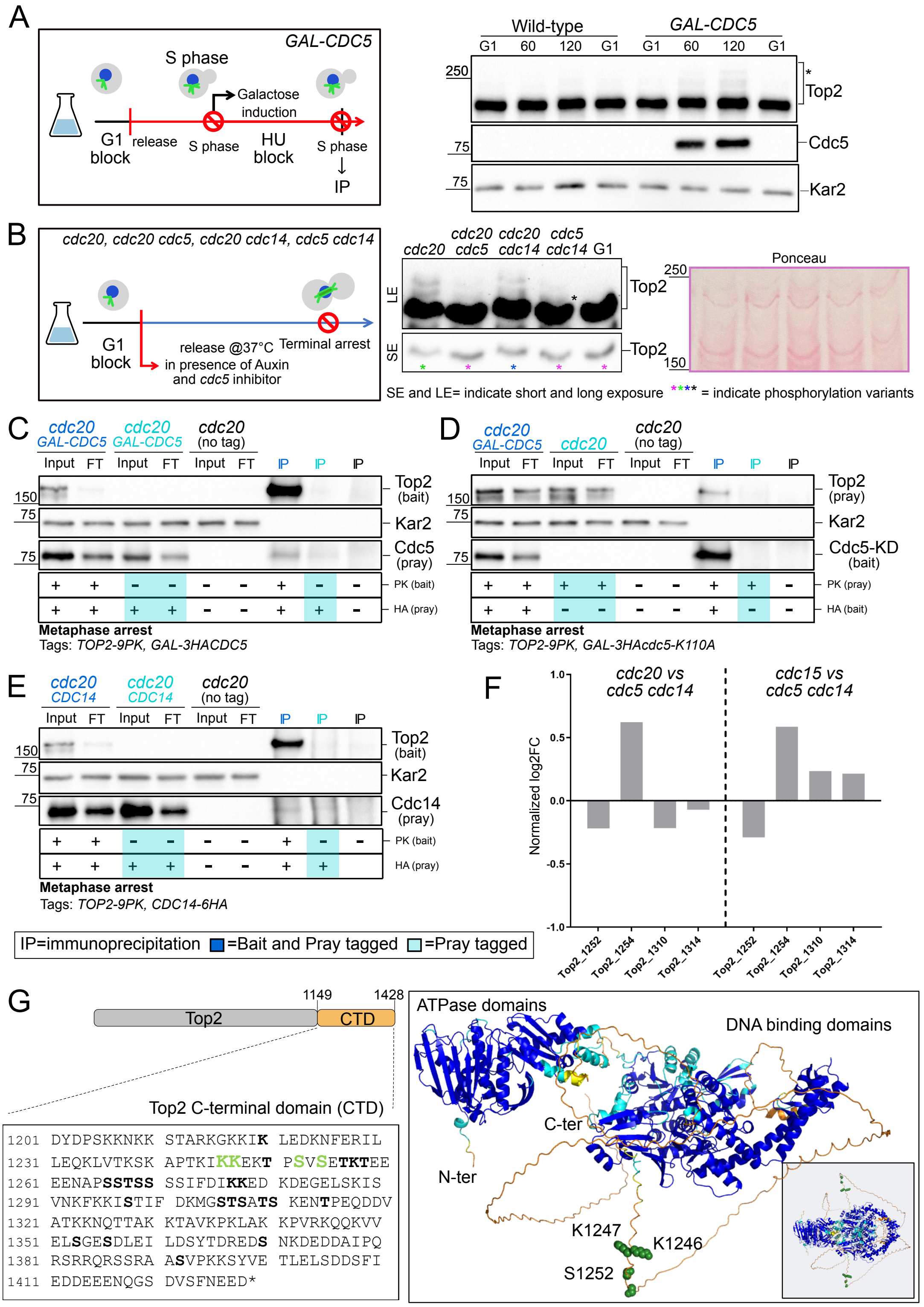


Supplementary Figure 11: **Cdc5 and Cdc14 affect Top2 phosphorylation.**
(**A**) *TOP2-9PK* (Ry7921) and *TOP2-9PK GAL-CDC5-3MYC* (Ry9325) cells were arrested in S phase with hydroxyurea in YPR medium for 3h. Upon arrest, galactose was added to

induce Cdc5 overexpression. Samples were collected at the indicated times to assess Top2 and Cdc5 levels and migration. Kar2 served as a loading control.
(**B**) *CDC20-AID (Ry8315), CDC20-AID cdc5-as1 (Ry8785), CDC20-AID cdc14-1 (Ry8782) and cdc5-as1 cdc14-1 (Ry7998)* cells carrying a *TOP2-9PK* fusion were synchronously released from a G1 block in conditions restrictive for the proteins of interest. 180 minutes after release, samples were collected for immunostaining against Top2 on Phos-tag SDS-PAGE.
(**C-D**) *CDC20-AID* (Ry4852), *CDC20-AID GAL-3HACDC5* (Ry11306) and *CDC20-AID GAL-3HACDC5 TOP2-9PK* (Ry11294) (**C**) or *CDC20-AID* (Ry4852), *CDC20-AID TOP2-9PK* (Ry8315) and *CDC20-AID GAL-3HAcdc5-K110A TOP2-9PK* (Ry11289) (**D**) cells were synchronously released from a G1 block into metaphase (*cdc20* arrest) in YPR medium. After arrest, galactose was added for 1.5 h to induce Cdc5 expression, after which samples were collected for Top2-PK immunoprecipitation and analyzed for interaction with Cdc5-HA (**C**) or for Cdc5-KD-HA immunoprecipitation and interaction with Top2-PK (**D**). Kar2 served as loading control.
(**E**) *CDC20-AID* (Ry4852), *CDC20-AID CDC14-6HA* (Ry11298) and *CDC20-AID CDC14-6HA TOP2-9PK* (Ry11279) were treated as in (**A**). At the terminal arrests, samples were collected for Top2-Pk immunoprecipitation and analyzed for interaction with Cdc14-HA. Kar2 served as loading control.
(**F**) The graph shows the phospho-residues associated with Top2 and their log2 fold changes (generated comparing *cdc14 cdc5* cells with either *cdc20* or *cdc15* cells, as indicated on the top of the graphs), normalized by the corresponding log2 fold change of Top2 total protein amount, to account for variations in protein abundance. Details of the experimental procedures and analyses are provided in (61). Briefly, *CDC20-AID* (Ry4852)*, cdc14-1 cdc5-as1* (Ry1602), and *cdc15-as1* (Ry1112) cells were arrested in G1 and subsequently released under conditions restrictive for the proteins of interest. Cells were harvested at their terminal arrest (~3 hours post-release) for protein extraction and processed for TMT mass spectrometry analysis, with three biological replicates for *CDC20-AID* and *cdc14-1 cdc5-as1* cells and two biological replicates for *cdc15-as1* cells. To identify phospho-residues with significant differences between mutants, a false discovery rate (FDR) of 0.001 (S0 at 0.05) was applied, along with an additional cutoff of log2 fold change < -0.5 or > 0.5. The log2 fold changes of the phospho-residues obtained through this analysis were also plotted against the corresponding protein log2 fold changes. For each phospho-residue, the log2 fold change was calculated by comparing *cdc14-1 cdc5-as1* with *CDC20-AID* or *cdc15-as1* cells.
(**G**) Left: Schematic representation and amino acid sequence of Top2 C-terminal domain. Serines (S1252, S1254) and lysines (K1246 and K1247) of interest are highlighted in green. Right: AlphaFold3-generated model of the functional dimeric yeast Top2 protein (small square) and enlarged view of the model, from which one Top2 monomer has been removed for clarity. The model is shown as cartoon and the colors represent the per-residue confidence scores (pLDDT), with dark blue indicating highest confidence and orange indicating lowest confidence/disordered regions. The side chains of serine (S1252 and S1254) and lysine (K1246 and K1247) residues of interest are highlighted as green spheres.

| Name | Genotype |
|---|---|
| Ry1 | *MATa, ade2-1, leu2-3, ura3, trp1-1, his3-11,15, can1-100, GAL, psi+* |
| Ry1112 | *MATa, cdc15::cdc15-as1(L99G)::URA3* |
| Ry1573 | *MATa, cdc14-1* |
| Ry1602 | *MATa, cdc14-1, cdc5L158G* |
| Ry2725 | *MATa, cdc14-1, cdc5L158G, PDS1-HA -LEU2::pds1, SCC1myc18::TRP1* |
| Ry2795 | *MATa, cdc14-1, cdc5L158G, scc1::HIS3, SCC1-TEV268-HA::LEU2, GAL-NLS-9MYC-TEV Protease-NLS::TRP1 (10 Iintegrants checked by southern)* |
| Ry2446 | *MATa, cdc5L158G* |
| Ry3203 | *MATa, cdc14-1, cdc5L158G, MET-CDC20::URA3* |
| Ry4852 | *MATa, ura3::pADH1-OsTIR1-9MYC::URA3, CDC20-aid::KanMX* |
| Ry4931 | *MATa, scc1::HIS3, SCC1-TEV268-HA::LEU2, pGAL-NLS-9MYC-TEV Protease-NLS::TRP1 (10 integrants), ura3::pADH1-OsTIR1-9MYC::URA3, CDC20-aid::KanMX* |
| Ry5156 | *MATa, cdc14-1, ura3::PGAL-NLS-NLS-ChVTOP2-PK3::URA3, cdc5L158G* |
| Ry5172 | *MATa, cdc14-1, cdc5-as1(L158G), ura3::pGAL-TOP2::URA3 (+)* |
| Ry5272 | *MATa, cdc14-1, cdc5-as1(L158G), YEN1ON-myc18::URA3* |
| Ry5815 | *MATa, cdc14-1, cdc5-as1(L158G), LEU2::TetR-GFP, ChrXV326bp::tetO112::KanMX* |
| Ry5824 | *MATa, cdc14-1, cdc5-as1(L158G), LEU2::TetR-GFP, ChrXVhis3::tetO112::HIS3* |
| Ry5956 | *MATa, cdc14-1, cdc5-as1(L158G), trp1::PGAL-CIN8::TRP1 (x1)* |
| Ry6099 | *MATa, cdc14-1, cdc5-as1(L158G), HTB2-Cherry::HIS3, cfi1::CFI1-GFP::KanMX6* |
| Ry6101 | *MATa, cdc14-1, HTB2-Cherry::HIS3, cfi1::CFI1-GFP::KanMX6* |
| Ry6105 | *MATa, HTB2-Cherry::HIS3, cfi1::CFI1-GFP::KanMX6* |
| Ry6289 | *MATa, HTB2-Cherry::HIS3, cfi1::CFI1-GFP::KanMX6, cdc5L158G* |
| Ry6589 | *MATa, cdc14-1, cdc5-as1(L158G), ndc10-1, HTB2-Cherry::HIS3, ura3::pAFS125-TUB1p-GFPTUB1::URA3* |
| Ry6590 | *MATa, ndc10-1, HTB2-Cherry::HIS3, ura3::pAFS125-TUB1p-GFPTUB1::URA3* |
| Ry6591 | *MATa, cdc14-1, cdc5-as1(L158G), HTB2-Cherry::HIS3, ura3::pAFS125-TUB1p GFPTUB1::URA3* |
| Ry7037 | *MATa, cdc14-1, smc2-8, cdc5-as1(L158G), ura3::pGAL-NLS- NLS-ChVTOP2-Pk3::URA3* |
| Ry7098 | *MATa, cdc14-1, cdc5-as1(L158G), LEU2::TetR-GFP, ChrXV1070K::tetO112::Hph, ura3::pRS306-mCherry- TUB1::URA3* |
| Ry7220 | *MATa, cdc14-1, cdc5-as1(L158G), YEN1ON-myc9::KanMX4 PGAL-CIN8::TRP1 (x1)* |
| Ry7411 | *MATa, ChrXV his3::tetO112::HIS3, LEU2::TetR-GFP, ura3::pADH1-OsTIR1-9MYC::URA3, CDC20-aid::KanMX* |
| Ry7481 | *MATa, LEU2::TetR-GFP, ChrXV 1070K::tetO112::Hph ura3::pADH1-OsTIR1-9MYC::URA3, CDC20-aid::KanMX* |
| Ry7519 | *MATa, LEU2::TetR-GFP, ChrXV 326K::tetO112::KanMX,* |

| | |
|---|---|
| Ry7563 | *MATa, scc1::HIS3, SCC1-TEV268-HA::LEU2, GAL-NLS-9MYC-TEV Protease-NLS::TRP1 (10 lintegrants), ura3::pADH1-OsTIR1-9MYC::URA3, CDC20-aid::KanMX, cdc15::CDC15-as1(L99G)::URA3* |
| Ry7874 | *MATa, leu2::pTEF1-osTIR::LEU2, CDC20-aid::KanMX.* |
| Ry7921 | *MATa, TOP2-PK9::HIS3* |
| Ry7998 | *MATa, TOP2-PK9::HIS3, cdc14-1, cdc5L158G* |
| Ry8315 | *MATa, TOP2-PK9::HIS3, ura3::pADH1-OsTIR1-9MYC::URA3, CDC20-aid::KanMX* |
| Ry8782 | *MATa, cdc14-1, TOP2-PK9::HIS3, ura3::pADH1-OsTIR1- 9MYC::URA3, CDC20-aid::KanMX* |
| Ry8785 | *MATa, TOP2-PK9::HIS3, cdc5L158G, ura3::pADH1-OsTIR1- 9MYC::URA3, CDC20-aid::KanMX* |
| Ry9325 | *MATa, ura::PGAL-3Myc-CDC5::URA3, TOP2-PK9::HIS3* |
| Ry10270 | *MATa, cdc14-1, leu2::PGAL-3HA-YEN1::LEU2, cdc5L158G* |
| Ry10473 | *MATa, cdc14-1, TOP2-HA3x-kanMX:pCC117* |
| Ry10485 | *MATa, cdc14-1, top2-SNM-HA3x-kanMX:pML251* |
| Ry10521 | *MATa, TOP2-PK9::HIS3, ura3::pADH1-OsTIR1- 9MYC::URA3, CDC20-aid::KanMX, cdc15::CDC15-as1(L99G)::URA3* |
| Ry10532 | *MATa, cdc15::CDC15-as1(L99G)::URA3***,** *trp::pGAL-ULP2:TRP* |
| Ry10539 | *MATa, cdc15::CDC15-as1(L99G)::URA3***,***TOP2-HA3x-kanMX:pCC117* |
| Ry10542 | *MATa, cdc15::CDC15-as1(L99G)::URA3***,** *top2-SNM-HA3x-kanMX:pML251* |
| Ry11225 | *MATa, leu2::pTEF1-osTIR::LEU2, CDC20-aid::KanMX. ura3::pGAL-NLS-NLS-ChVTOP2-Pk3::URA3* |
| Ry11241 | *MATa, TOP2-PK9::HIS3, leu2::pTEF1-osTIR::LEU2, CDC20-aid::KanMX, 3xGAL-CDC14::URA3, cdc5L158G* |
| Ry11247 | *MATa, cdc14-1,cdc5L158G, LEU2::TetR-GFP, ChrXV 326K::tetO112::Kan, Spc42-mScarlet-I::LEU2* |
| Ry11273 | *MATa, LEU2::TetR-GFP, ChrXV 326K::tetO112::Kan, ura3::pADH1-OsTIR1-9MYC::URA3, CDC20-aid::KanMX, Spc42-mScarlet-I::LEU2* |
| Ry11279 | *MATa, TOP2-PK9::HIS3, ura3::pADH1-OsTIR1-9MYC::URA3, CDC20-aid::KanMX, CDC14-6HA::kanMX6* |
| RY11289 | *MATa, leu2::pTEF1-osTIR::LEU2, TOP2-PK9::HIS3, CDC20-aid::KanMX, ura3::1xpGAL-HA3cdc5-K110A::URA3* |
| Ry11294 | *MATa, leu2::pTEF1-osTIR::LEU2, TOP2-PK9::HIS3, CDC20-aid::KanMX, ura3::1xpGAL-HA3CDC5::URA3* |
| Ry11298 | *MATa, ura3::pADH1-OsTIR1-9MYC::URA3, CDC20-aid::KanMX, CDC14-6HA::kanMX6.* |
| Ry11306 | *MATa, leu2::pTEF1-osTIR::LEU2, cdc5L158G, CDC20-aid::KanMX, ura3::1xpGAL-HA3CDC5::URA3* |
| Ry11307 | *MATa, TOP2-PK9::HIS3, leu2::pTEF1-osTIR::LEU2, CDC20-aid::KanMX, 3xGAL-CDC14::URA3, cdc14-1* |

Table S1: **Yeast strains used in this study**

| Chromosome | Centromere coordinates | |
|---|---|---|
| chrI | 151523 | 151524 |
| chrII | 238265 | 238266 |
| chrIII | 114443 | 114444 |
| chrIV | 448766 | 448767 |
| chrV | 152046 | 152047 |
| chrVI | 148569 | 148570 |
| chrVII | 496979 | 496980 |
| chrVIII | 105645 | 105646 |
| chrIX | 355687 | 355688 |
| chrX | 436366 | 436367 |
| chrXI | 440188 | 440189 |
| chrXII | 150888 | 150889 |
| chrXIII | 268090 | 268091 |
| chrXIV | 628817 | 628818 |
| chrXV | 326643 | 326644 |
| chrXVI | 556015 | 556016 |

Supplementary Table 2: **Centromere coordinates**

| comparison | statistics | p_value | stars | Statistical test |
|---|---|---|---|---|
| Top2 MFI (a.u.) | 865,1112029 | 6,033E-186 | **** | kruskal |
| Top2 MFI (a.u.) - cdc20 + NOCO vs cdc20 | 129986 | 5,58842E-86 | **** | mannwhitneyu |
| Top2 MFI (a.u.) - cdc20 + NOCO vs cdc5 cdc14 | 232587 | 1,9516E-117 | **** | mannwhitneyu |
| Top2 MFI (a.u.) - cdc20 + NOCO vs cdc20 cdc5 | 176124 | 1,1923E-105 | **** | mannwhitneyu |
| Top2 MFI (a.u.) - cdc20 + NOCO vs cdc20 cdc14 | 68746 | 7,35853E-62 | **** | mannwhitneyu |
| Top2 MFI (a.u.) - cdc20 vs cdc5 cdc14 | 71784 | 0,149051555 | ns | mannwhitneyu |
| Top2 MFI (a.u.) - cdc20 vs cdc20 cdc5 | 50137 | 0,858429686 | ns | mannwhitneyu |
| Top2 MFI (a.u.) - cdc20 vs cdc20 cdc14 | 29303 | 1,51562E-18 | **** | mannwhitneyu |
| Top2 MFI (a.u.) - cdc5 cdc14 vs cdc20 cdc5 | 83378 | 0,036113663 | * | mannwhitneyu |
| Top2 MFI (a.u.) - cdc5 cdc14 vs cdc20 cdc14 | 51296 | 1,49454E-18 | **** | mannwhitneyu |
| Top2 MFI (a.u.) - cdc20 cdc5 vs cdc20 cdc14 | 41225 | 2,76851E-25 | **** | mannwhitneyu |

Supplementary Table 3: **Mean nuclear Top2 fluorescence intensity**

| comparison | statistics | p_value | stars | Statistical test |
|---|---|---|---|---|
| Top2 CV | 296,0973672 | 7,52665E-63 | **** | kruskal |
| Top2 CV - cdc20 + NOCO vs cdc20 | 101325 | 1,64771E-24 | **** | mannwhitneyu |
| Top2 CV - cdc20 + NOCO vs cdc5 cdc14 | 193934 | 1,39842E-48 | **** | mannwhitneyu |
| Top2 CV - cdc20 + NOCO vs cdc20 cdc5 | 146276 | 1,61234E-43 | **** | mannwhitneyu |
| Top2 CV - cdc20 + NOCO vs cdc20 cdc14 | 53872 | 1,09922E-19 | **** | mannwhitneyu |
| Top2 CV - cdc20 vs cdc5 cdc14 | 74606 | 0,016276218 | * | mannwhitneyu |
| Top2 CV - cdc20 vs cdc20 cdc5 | 55381 | 0,03822463 | * | mannwhitneyu |
| Top2 CV - cdc20 vs cdc20 cdc14 | 19977 | 0,489230504 | ns | mannwhitneyu |
| Top2 CV - cdc5 cdc14 vs cdc20 cdc5 | 89715 | 0,731107516 | ns | mannwhitneyu |
| Top2 CV - cdc5 cdc14 vs cdc20 cdc14 | 32098 | 0,206649348 | ns | mannwhitneyu |
| Top2 CV - cdc20 cdc5 vs cdc20 cdc14 | 24325 | 0,309933723 | ns | mannwhitneyu |

Supplementary Table 4: **Coefficient of variation**
